# A Paradigm For Calling Sequence In Families: The Long Life Family Study

**DOI:** 10.1101/2024.05.23.595584

**Authors:** E. Warwick Daw, Jason A. Anema, Karen Schwander, Shiow Jiuan Lin, Lihua Wang, Mary Wojczynski, Bharat Thyagarajan, Nathan Stitziel, Michael A. Province

## Abstract

Over Several years, we have developed a system for assuring the quality of whole genome sequence (WGS) data in the LLFS families. We have focused on providing data to identify germline genetic variants with the aim of releasing as many variants on as many individuals as possible. We aim to assure the quality of the individual calls. The availability of family data has enabled us to use and validate some filters not commonly used in population-based studies. We developed slightly different procedures for the autosomal, X, Y, and Mitochondrial (MT) chromosomes. Some of these filters are specific to family data, but some can be used with any WGS data set. We also describe the procedure we use to construct linkage markers from the SNP sequence data and how we compute IBD values for use in linkage analysis.

## Introduction

The Long Life Family Study (LLFS) is a multi-center study designed to identify genetic contributions to healthy aging^1^. Data collection is handled at four field sites (Boston University, Columbia University, University of Pittsburgh, and University of Southern Denmark), and biospecimens are processed and stored at the University of Minnesota. Washington University houses the Data Management Coordinating Center for the study and genotyping as well as several other omics are done at Washington University. Starting in 2006, the study ascertained siblings in their 80’s, as well any spouses, children, and spouses of the children who were willing to participate^2^. In the most recent funding cycle, grandchildren of the original proband siblings were also collected. There have been three extensive in person examinations of the participants, and annual telephone follow-up interviews. Thus, the LLFS study provides a rich longitudinal data set to study healthy aging in families.

When we started getting next generation whole genome sequence data for LLFS, we set out to follow established procedure. We were aware that most studies doing sequencing were population rather than family studies, and that the relatedness of our subjects would allow us to carry out additional quality assurance measures that were not possible in studies of unrelated individuals. However, we were also told that while the processing and alignment of sequence data into CRAM files was well standardized^3^, there was room for interpretation in the actual calling of variants. As a result, we adopted a cautious approach, using established procedures but verifying the results with the tools available to us, including cross-checking the data within pedigrees. We have been constantly improving the methods we use for several years, and this manuscript describes our current sequence calling pipeline. Some of the procedures we use are only useful to studies with family data, but others may be useful to any study with sequence data. Accordingly, we describe our data processing steps here in full.

### Standard sequencing and alignment procedures

DNA samples were sent to the McDonnell Genome Institute (MGI) at Washington University in St. Louis for Whole genome sequencing (WGS) on Illumina Sequencers with 150bp reads. MGI performed alignment procedures. The steps of this workflow are listed in Table 1, including sequence alignment to build GRCh38 with BWA-MEM^4^, marking duplicates with Picard^5^, base quality score recalibration with Genome Analysis Toolkit (GATK)^6^, and lossless conversion to CRAM format with SAMtools^7^.

**Table 1.** Alignment Tools C Specifications from MGI.

| Process | Program | Version | Description and optional parameters |
| --- | --- | --- | --- |
| Align | BWA-MEM | 0.7.15 | Reference build38 (GRCh38DH) Opt: -K 10,000,000 -t -p -Y |
| Mark Duplicates | Picard | 2.4.1 |  |
| Sort | Sambamba | 0.6.4 |  |
| BQSR | GATK BaseRecalibrator | 3.6 | dbSNP 138, Known Indels, Mills and 1000G Indels<br>GATK hg38 Resource Bundle |
| Apply BQSR | GATK PrintReads | 3.6 |  |
| Convert to CRAM | SAMtools | ≥1.3.1 |  |

The steps of quality control performed at MGI (listed in Table 2) included generation of library insert size, alignment and aggregate target/bait interval coverage metrics as well as per target and per base coverage metrics with Picard. Inputs to CollectWgsMetrics are intervals for the autosomal chromosomes to evaluate coverage. CollectGcBiasMetrics was run to calculate G+C bias metrics across bins of the reference genome. SAMtools v1.3.1 flagstat was run to additionally summarize alignment metrics. Finally, FREEMIX values were generated using the VerifyBamID software to identify contamination based on known SNPs^8^.

**Table 2.** Quality Control Tools C Specifications from MGI.

| Process | Program | Version | Description and optional parameters |
| --- | --- | --- | --- |
| CollectInsertSizeMetrics | Picard | 2.14.0 |  |
| CollectAlignmentSummaryMetrics | Picard | 2.14.0 |  |
| CollectWgsMetrics | Picard | 2.14.0 | Autosomal chromosome intervals, |
| CollectGcBiasMetrics | Picard | 2.14.0 |  |
| Flagstat | SAMtools | 1.3.1 |  |
| FREEMIX | verifyBamID | 1.1.3 | Omni 2.5M SNP from 1000G<br>verifyBamID and Omni VCF Download |

### Standard Variant Calling

Variant calling was performed at the LLFS Data Management Coordinating Center, Division of Statistical Genomics at Washington University in St. Louis, using GATK4.1.0.0. GATK HaplotypeCaller^9^ was used to call variants from the CRAM files and create subject-level GVCF files. Autosomal chromosomes were called as diploid, while chromosomes Y (on male subjects only) and MT were called as haploid. For chromosome X, female subjects were called as diploid, along with males for variants in the pseudo-autosomal regions (PAR). Male subjects in non-PAR regions on chromosome X were called as haploid. For males, variants in the X-transposed region (XTR) were found to be of particularly problematic quality, so we have not released the data for the XTR in males.

Next, these files were combined using GATK CombineGVCFs, and jointly called using GATK GenotypeGVCFs (GATK can appropriately handle VCF files with different ploidies). Finally, diallelic variants were extracted using GATK SelectVariants. Our processing for other variant types is slightly different. The vast majority of variants are diallelic, so the remainder of this document will focus on our processing to them.

In order to facilitate processing and reduce the size of individual files, autosomal chromosomes were divided into 260 12.5mb “chunks.” After QC, three chunks on the p-arms of acrocentric chromosomes (13,14,15) were completely void of quality called variants.

Because variant calling at the end of a chunk can be problematic due to reads split between chunks, a 10kb region was added to the ends of each chunk, so that it overlaps with the surrounding chunks. For example, the chr1:12500001-25000000 chunk was actually called as 12,490,000-25,010,000. After the variant calling was completed, the overlapping 10kb regions were removed from each chunk, leaving the original 260 non-overlapping chunks spanning the autosome. Chromosomes X and Y were similarly called in chunks, but Mt is sufficiently small that it was called as a whole.

### Sample Verification

To verify familial relationships and identify any accidental sample switches, we first compared each subject’s WGS data with their cleaned GWAS chip data; any individual that did not match >99% was set aside for further review.

Next, we used KING (Kinship-based INference for Gwas)^10^ to verify relationships *within* pedigrees and to look for relationships *between* pedigrees. This technique allowed us to identify and correct misreported relationships within pedigrees (mostly half-siblings reported as full siblings), and also enabled us to identify accidental sample switches. In a few cases, these switches occurred between individuals from 2 completely different families that visited the field center on the same day, where blood samples were likely mislabeled.

### Tools for evaluating filters

In evaluating where to set our filters, we used several metrics. The Mendel error rate (a mismatch at a marker among family members) was one metric we used in evaluating our filters. The heterozygosity rate can also be used: a low rate for a subject can indicate a missing haplotype, while a high rate can indicate contamination. Another metric available in family data is looking at the excess recombination rate when haplotyping the variant calls: even if genotyping errors do not cause Mendel errors, they can cause tight double recombination events to be falsely inferred when haplotyping the data.

### Initial Filters

MGI provided CRAM files for all failed samples that received sufficient processing to generate a CRAM file (at our request), so we could review the MGI process for identifying failures. The two primary reasons for failure were evidence of contamination, as measured by the FREEMIX statistic, and insufficient haploid coverage (HC). We noted that although MGI called HC<30 (i.e. below the target coverage) as a failure, many of these seemed to provide adequate sequence calls if there were no secondary issues. In order to save as many WGS samples as possible, we elected to only automatically fail samples with HC<20, although some of these subjects with 20<=HC<30 were removed due to failing later filters. Additionally, MGI called FREEMIX>0.05 as a failure, but we found this to be too liberal; as a result, we have filtered subjects with FREEMIX>0.03, the threshold suggested in the original paper^8^. So, our initial filter was to exclude subjects with HC<20 and FREEMIX>0.03. We made note of samples with HC between 20 and 30 so that researchers could use them at their discretion.

Note that one reason for a HC filter is to reduce the chance of an erroneous call due to only one of the parental haplotypes being present. At HC=20, there is a 1 in 2^20^ (1,048,576) chance that all the reads are from the mom and the same chance that all are from the dad. Such a situation will only lead to a genotyping error if the individual is heterozygous (got different variants from the mom and dad), and the chance of that is at most 50% with a diallelic variant. The calling algorithm is a little more complicated than just looking at the alleles present, but the point here is that with HC>=20, the chance of a miscall due to only one parental haplotype being present is on the order of one in a million or less. While one might be tempted to accept a lower HC threshold, we found that doing so greatly increased both the Mendel error rate and the excess recombination rates. At HC=10, the chance of an error is more on the order of 1 in 1000, which may not sound that bad, but a small amount of data with an error rate on that order can cause a 100-fold increase in the number of recombinations found in Lander-Green haplotyping. Thus, a cut off of HC=20 seems justified.

After these initial filters, we found a small number of samples with large portions of the genome with less than 20x coverage (PCT_20x<0.8) to also be problematic, and these were also excluded from the primary analysis files.

These initial filters are roughly similar to filters used by the TOPMed sequencing centers, and indeed, for many groups, the quality control process would end at this step. However, we have taken further steps to produce very high quality data for analysis, including some steps that are only available with family data.

### Autosomal Chromosome Processing

There are slight differences in the Autosomal and X, Y, and MT chromosome processing, so we will first describe the Autosomal (1-22) chromosome processing for SNPs and INDELs.

### Individual Call Depth Filter

For each autosomal sequence call at each variant site, the individual call was set to missing for depth less than 20 (at least twenty sequence reads covering the variant site) or greater than 300. This filter was selected to balance the competing needs of quality data and complete data. With low depth, one risk is that both haplotypes are not represented, while with extremely high depth, there is high risk that the variant is an artifact (e.g., a site with high homology across the genome generating a false variant due to slight differences between the different homologous sites). Individual calls that do not meet these basic filter criteria were set to missing.

After setting individual variant calls to missing based on the depth filters, there were some variants that became either monomorphic or completely missing. We removed these non-informative variants from the VCF files, since there is little benefit to keeping them.

### Excess Heterozygosity Filter (whole variant filter)

In our pilot sequence data, we observed some common variants with very high heterozygosity. While such variants with excess heterozygosity may have high PHRED quality scores, they usually fail a Hardy-Weinberg Equilibrium test (HWE). We examined several of these variants further. BLAST searches for the sequence reads containing these variants had multiple hits across the genome. Examination of the sequence reads around these sites found that there were more than two haplotypes present, suggesting that reads from multiple homologous regions were being mapped to the same location. Also, in some of these regions, multiple SNPs were called in short regions, and the depth of coverage tended to be higher than the target. This evidence suggests that the excess heterozygosity markers are artifacts as a result of different regions on different chromosomes having similar sequence. Consequently, we chose to filter out variants which failed the HWE test because of excess heterozygosity with p-value <1e-6, which removed most of these markers. Note that a diallelic marker has excess heterozygosity if the proportion of heterozygotes is >2pq where p and q are the frequencies of the 2 alleles. There were still some markers with heterozygosity > 0.55, and we subsequently removed those as well. Figures 1a and 1b show SNP heterozygosity by HWE p-value, before and after the excess heterozygosity filter.

We did not filter markers failing the HWE test because of excess homozygosity since such failure can be due to population substructure or due to the marker being under selective pressure. However, such failure can alternatively be an indication of a problematic marker, thus these markers should be used with appropriate caution. Indeed, most of the remaining markers with excess homozygosity are removed after the call rate filter described next. INDEL heterozygosity figures look similar and are not shown here.

**Figures 1a-1c.**
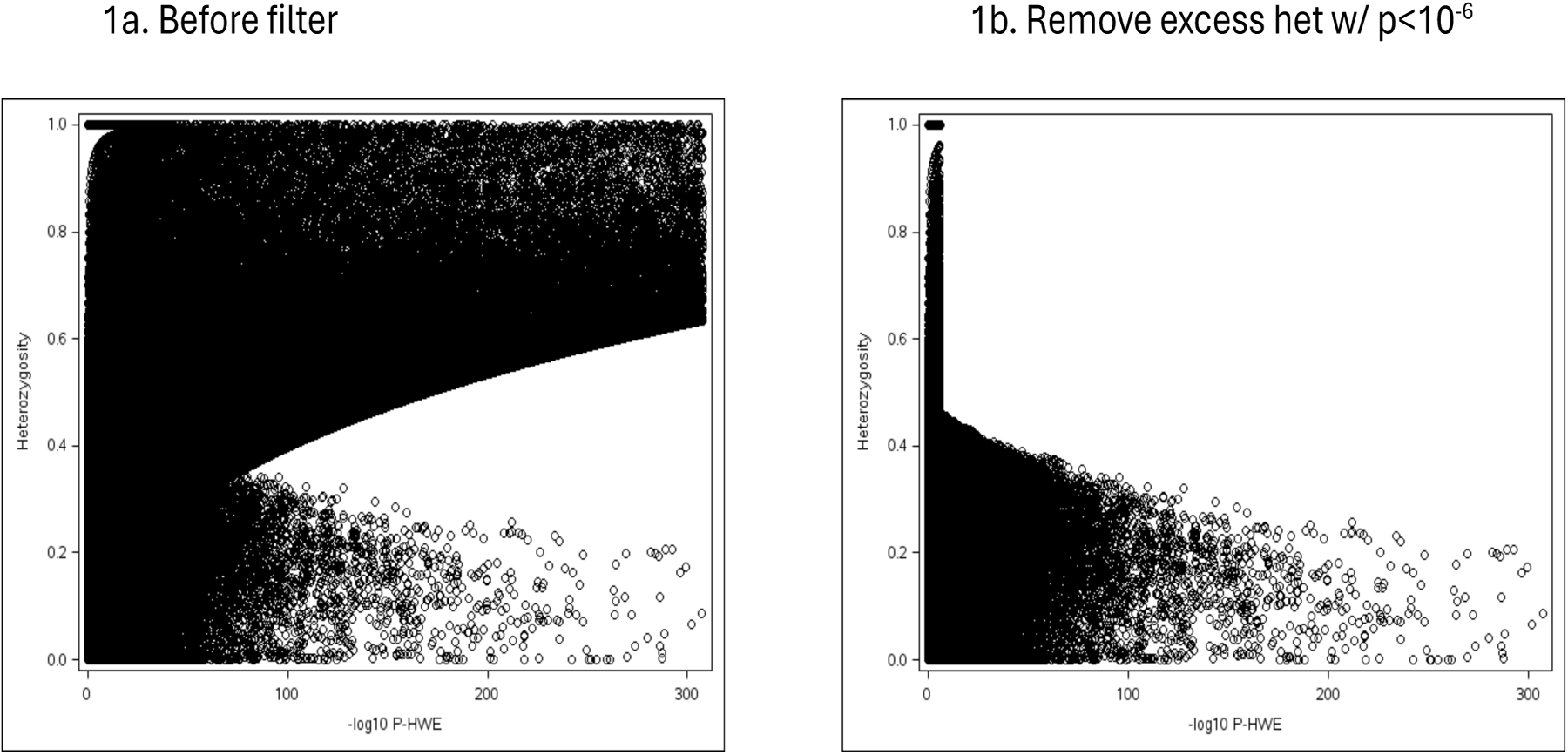
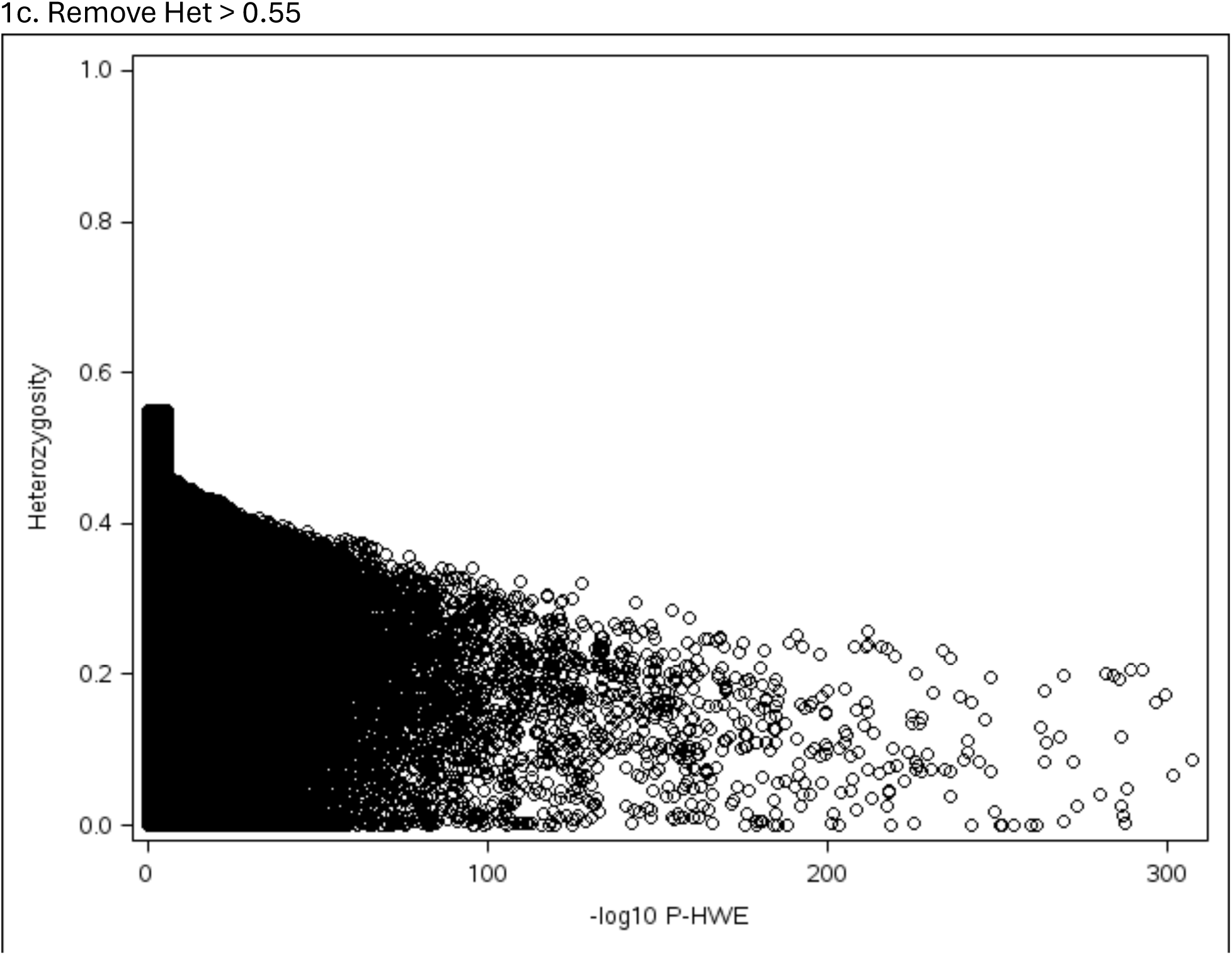
Before and After Excess Heterozygosity Filter for SNPs

### Variant Call Rate Filter

Low call rate can be indicative of poor variant quality or other issues. Variants with many lower depth calls now have higher missing rates, since calls with depth<20 were set to missing. After reviewing histograms and summary statistics, we filtered variants with a call rate of less than 90%. This allowed us to remove extreme outliers, while still retaining approximately 93% of the variants.

### Summary of Preliminary Filter effects

The effects of the filters to this point can be seen in Table 3a for SNPs and 3b for INDELs.

**Table 3a.** Preliminary QC Statistics on Autosomal Diallelic SNPs in VCF Files.

| Chr | # Of<br>Chunk<br>s | #SNPs<br>before QC | % calls set<br>to missing<br>(depth<20) | # (Additional) SNPs Dropped |  |  |  |  |  | After<br>Preliminar<br>y QC |
| --- | --- | --- | --- | --- | --- | --- | --- | --- | --- | --- |
|  |  |  |  | All Missing or<br>Monomorphic |  | HETZ > .55 OR<br>Excess HETZ p<<br>1E-6 |  | Call Rate <0.90 |  |  |
|  |  |  |  | N | % | N | % | N | % |  |
| 1 | 20 | 6,041,788 | 8.92 | 882,984 | 14.61 | 34,280 | 0.57 | 506,428 | 8.38 | 4,618,096 |
| 2 | 20 | 6,031,518 | 6.44 | 678,976 | 11.26 | 25,341 | 0.42 | 333,838 | 5.53 | 4,993,363 |
| 3 | 16 | 4,919,431 | 5.79 | 486,454 | 9.89 | 20,190 | 0.41 | 251,791 | 5.12 | 4,160,996 |
| 4 | 16 | 4,739,226 | 5.73 | 475,902 | 10.04 | 14,366 | 0.30 | 231,549 | 4.89 | 4,017,409 |
| 5 | 15 | 4,411,137 | 6.05 | 445,056 | 10.09 | 13,469 | 0.31 | 227,797 | 5.16 | 3,724,815 |
| 6 | 14 | 4,146,828 | 5.23 | 378,573 | 9.13 | 14,973 | 0.36 | 190,127 | 4.58 | 3,563,155 |
| 7 | 13 | 4,039,040 | 5.85 | 441,726 | 10.94 | 27,648 | 0.68 | 215,235 | 5.33 | 3,354,431 |
| 8 | 12 | 3,755,922 | 5.64 | 352,335 | 9.38 | 10,003 | 0.27 | 178,059 | 4.74 | 3,215,525 |
| 9 | 12 | 3,297,380 | 8.07 | 461,811 | 14.01 | 38,497 | 1.17 | 239,503 | 7.26 | 2,557,569 |
| 10 | 11 | 3,435,318 | 5.82 | 380,978 | 11.09 | 19,872 | 0.58 | 196,941 | 5.73 | 2,837,527 |
| 11 | 11 | 3,605,103 | 6.14 | 545,963 | 15.14 | 12,430 | 0.34 | 205,086 | 5.69 | 2,841,624 |
| 12 | 11 | 3,404,503 | 6.34 | 456,625 | 13.41 | 12,671 | 0.37 | 200,632 | 5.89 | 2,734,575 |
| 13 | 9* | 2,687,943 | 9.94 | 412,009 | 15.33 | 17,491 | 0.65 | 246,980 | 9.19 | 2,011,463 |
| 14 | 8* | 2,174,828 | 5.62 | 190,457 | 8.76 | 12,342 | 0.57 | 124,445 | 5.72 | 1,847,584 |
| 15 | 8* | 2,077,853 | 6.75 | 229,201 | 11.03 | 17,057 | 0.82 | 139,445 | 6.71 | 1,692,150 |
| 16 | 8 | 2,319,750 | 6.31 | 240,557 | 10.37 | 15,492 | 0.67 | 139,594 | 6.02 | 1,924,107 |
| 17 | 7 | 2,090,856 | 5.75 | 265,403 | 12.69 | 19,116 | 0.91 | 123,916 | 5.93 | 1,682,421 |
| 18 | 7 | 2,104,383 | 8.65 | 362,214 | 17.21 | 6,864 | 0.33 | 139,383 | 6.62 | 1,595,922 |
| 19 | 5 | 1,572,583 | 4.47 | 146,084 | 9.29 | 10,691 | 0.68 | 89,562 | 5.70 | 1,326,246 |
| 20 | 6 | 1,707,822 | 5.70 | 204,934 | 12.00 | 36,171 | 2.12 | 100,115 | 5.86 | 1,366,602 |
| 21 | 4 | 1,030,725 | 7.08 | 95,879 | 9.30 | 27,341 | 2.65 | 76,068 | 7.38 | 831,437 |
| 22 | 5 | 1,093,076 | 7.51 | 109,701 | 10.04 | 30,900 | 2.83 | 90,698 | 8.30 | 861,777 |
| All | 238 | 70,687,013 | 6.54 | 8,243,822 | 11.66 | 437,205 | 0.62 | 4,247,192 | 6.01 | 57,758,794 |

**Table 3b.** Preliminary QC Statistics on Autosomal Diallelic INDELs in VCF Files.

| Chr | # Of<br>Chunk<br>s | #INDELs<br>before<br>QC | % calls set<br>to missing<br>(depth<20) | # (Additional) INDELs Dropped |  |  |  |  |  | After<br>Prelimina<br>ry QC |
| --- | --- | --- | --- | --- | --- | --- | --- | --- | --- | --- |
|  |  |  |  | All Missing or<br>Monomorphic |  | HETZ > .55 OR<br>Excess HETZ p<<br>1E-6 |  | Call Rate <0.90 |  |  |
|  |  |  |  | N | % | N | % | N | % |  |
| 1 | 20 | 580,285 | 8.33 | 108,048 | 18.62 | 1,702 | 0.29 | 48,020 | 8.28 | 422,515 |
| 2 | 20 | 603,181 | 7.28 | 103,350 | 17.13 | 1,710 | 0.28 | 41,424 | 6.87 | 456,697 |
| 3 | 16 | 496,008 | 6.91 | 82,764 | 16.69 | 1,211 | 0.24 | 33,065 | 6.67 | 378,968 |
| 4 | 16 | 485,762 | 7.19 | 80,873 | 16.65 | 854 | 0.18 | 33,437 | 6.88 | 370,598 |
| 5 | 15 | 445,378 | 7.34 | 76,047 | 17.07 | 695 | 0.16 | 30,249 | 6.79 | 338,387 |
| 6 | 14 | 439,165 | 6.98 | 71,342 | 16.24 | 1,375 | 0.31 | 29,939 | 6.82 | 336,509 |
| 7 | 13 | 406,788 | 7.58 | 69,306 | 17.04 | 2,016 | 0.50 | 30,661 | 7.54 | 304,805 |
| 8 | 12 | 358,385 | 7.07 | 60,355 | 16.84 | 595 | 0.17 | 23,834 | 6.65 | 273,601 |
| 9 | 12 | 306,817 | 8.79 | 57,037 | 18.59 | 2,391 | 0.78 | 26,064 | 8.49 | 221,325 |
| 10 | 11 | 338,144 | 7.25 | 57,907 | 17.12 | 1,297 | 0.38 | 25,274 | 7.47 | 253,666 |
| 11 | 11 | 334,278 | 7.06 | 59,053 | 17.67 | 622 | 0.19 | 23,744 | 7.10 | 250,859 |
| 12 | 11 | 340,500 | 7.31 | 59,701 | 17.53 | 909 | 0.27 | 24,765 | 7.27 | 255,125 |
| 13 | 9* | 264,328 | 8.51 | 49,179 | 18.61 | 763 | 0.29 | 21,252 | 8.04 | 193,134 |
| 14 | 8* | 225,926 | 7.35 | 36,832 | 16.30 | 804 | 0.36 | 17,594 | 7.79 | 170,696 |
| 15 | 8* | 212,362 | 8.34 | 37,117 | 17.48 | 1,303 | 0.61 | 18,128 | 8.54 | 155,814 |
| 16 | 8 | 206,809 | 8.38 | 36,304 | 17.55 | 934 | 0.45 | 18,418 | 8.91 | 151,153 |
| 17 | 7 | 212,772 | 7.23 | 37,454 | 17.60 | 1,336 | 0.63 | 17,036 | 8.01 | 156,946 |
| 18 | 7 | 199,878 | 7.63 | 37,491 | 18.76 | 405 | 0.20 | 13,758 | 6.88 | 148,224 |
| 19 | 5 | 162,095 | 7.32 | 24,721 | 15.25 | 860 | 0.53 | 15,086 | 9.31 | 121,428 |
| 20 | 6 | 161,728 | 6.79 | 28,115 | 17.38 | 2,160 | 1.34 | 11,912 | 7.37 | 119,541 |
| 21 | 4 | 103,521 | 8.50 | 17,736 | 17.13 | 1,664 | 1.61 | 9,201 | 8.89 | 74,920 |
| 22 | 5 | 104,478 | 8.71 | 17,803 | 17.04 | 1,887 | 1.81 | 10,146 | 9.71 | 74,642 |
| All | 238 | 6,988,588 | 7.53 | 1,208,535 | 17.29 | 27,493 | 0.39 | 523,007 | 7.48 | 5,229,553 |

### No “Blacklist” Used

There are “blacklists” of regions considered too difficult to sequence. Often all markers in these regions are filtered out^11–13^. These lists are evolving over time, but we have found that many of the markers in these regions are removed by our other filters (including some by the Mendel check described below). As we noted above, three of our processing “chunks” in p-arms of acrocentric chromosomes wound up with no calls after filtering, and sections of several others have no calls. Consequently, we did not apply a “blacklist” filter to this release ourselves, leaving the choice to use such a list open to individual researchers.

### B-allele Frequency checks for individuals (chromosome by chromosome)

We instituted checks on the B-allele frequencies (BAF) of each subject for each chromosome. B-allele frequency for a variant is the fraction of reads supporting a non-reference allele at that variant. In the early releases, we filtered subjects that appeared to have high Mendel error rates. It is necessary to remove all Mendel errors for most linkage analyses, and additionally large numbers of Mendel errors in a subject can indicate a problem with the sample. However, detecting Mendel errors requires information on relatives, and in the case of diallelic markers, only errors between parents and children can be detected. Furthermore, the amount of information to detect Mendel errors can differ vastly depending on how much relative information is available: in particular, in nuclear families with WGS data on both parents, many more Mendel errors can be detected than in nuclear families with one parent. We have found that the subjects with high Mendel errors also typically have unusual B-allele frequencies, and the B-allele check can be performed on individuals with no parents as well. Normally, the distribution of the B-allele frequency for diallelic markers will have three disjoint unimodal distributions, one for the reference homozygote (A/A), one for the refence/alternate homozygote (A/B), and one for the alternate homozygote (B/B). So, when performing subject-wide filtering, we now do B-allele frequency checks (for markers we retain Mendel error filtering described below).

In examining BAF, analysis of the A/B genotype band (in blue in figure 2 below) proved to be most useful. High A/B band variance (which can be caused by low depth of coverage, sample contamination, or chromosomal abnormalities), can result in the A/B band blending into the B/B and A/A bands (red and green in figure 2). This produces bad genotype calls. Samples with low genome wide coverage have high variance in the A/B band across the genome. We removed samples (mentioned previously) that did not have at least 80% of the genome covered at a depth of 20x. This removed all the samples with high A/B band variance across the genome.

In addition, we found that some subjects had unusual A/B bands only for specific chromosomes. These subjects have typical BAF for most of the genome, but an aberration on a specific chromosome, leading to bad SNP calls there. It should be emphasized that these phenomena are real and of biological significance. However, they also lead to bad germline SNPs calls. Thus, standard analyses may give erroneous results in the regions of these aberrations, so we decided to remove the SNP data for the chromosomes containing them. However, we only removed the chromosome if the aberration was severe enough to cause bad calls (see figure 3). Abnormal chromosomes were identified with a combination of A/B band variance and a Gaussian mixture model decomposition. The decision to keep or drop a chromosome was based on the presence of calls in an exclusion zone: In figures 2 and 3a, there is clear separation between the A/A, A/B, and B/B calls. However, in figure 3b, there is overlap between the A/A and A/B calls, and between the A/B and B/B calls.

BAF analysis identified a subject with multiple chromosomal abnormalities due to active leukemia at time of blood draw. We dropped this subject from the main analysis file.

**Figure 2.**
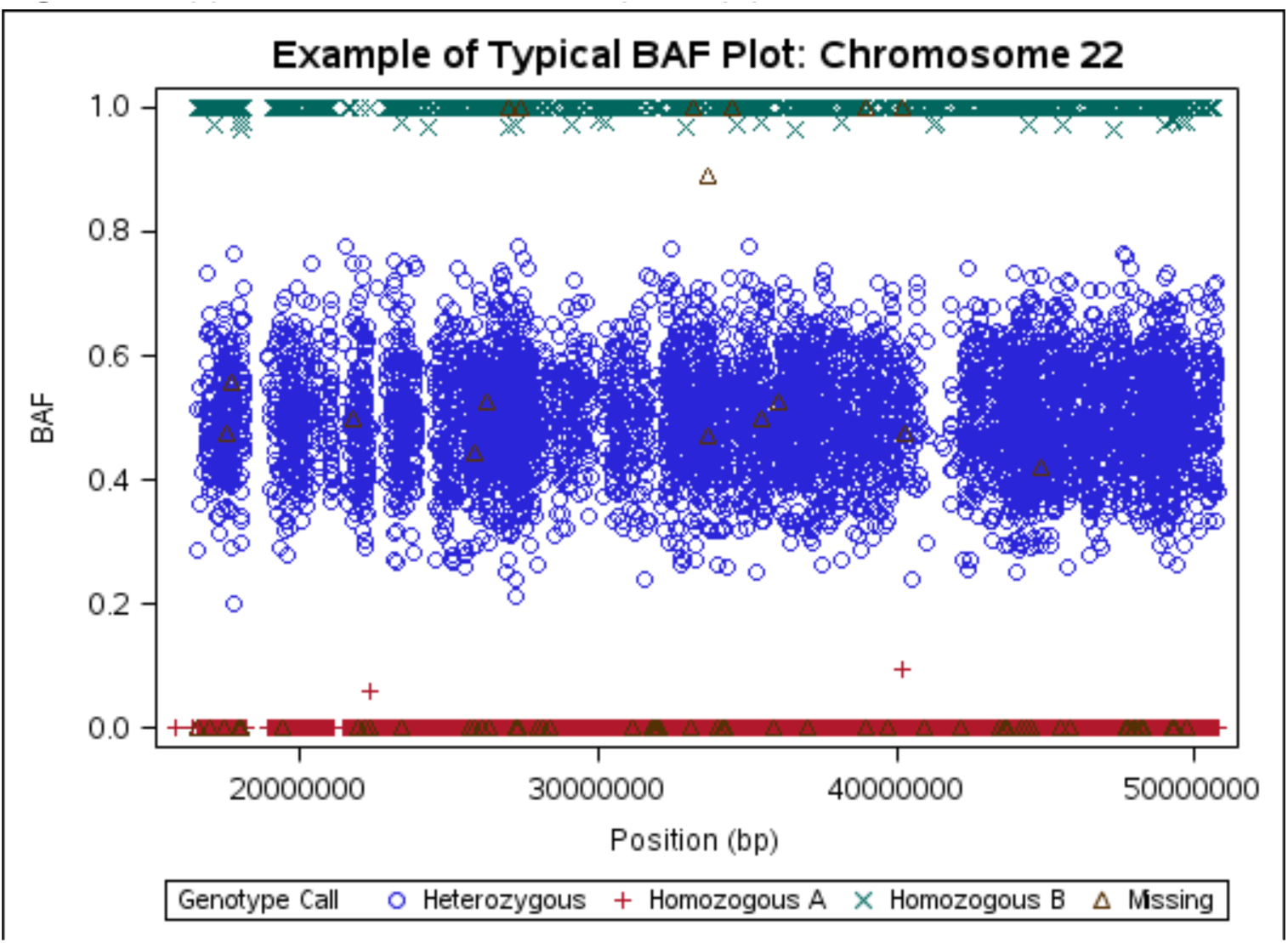
Typical WGS B-allele frequency plot

**Figure 3(a-b).**
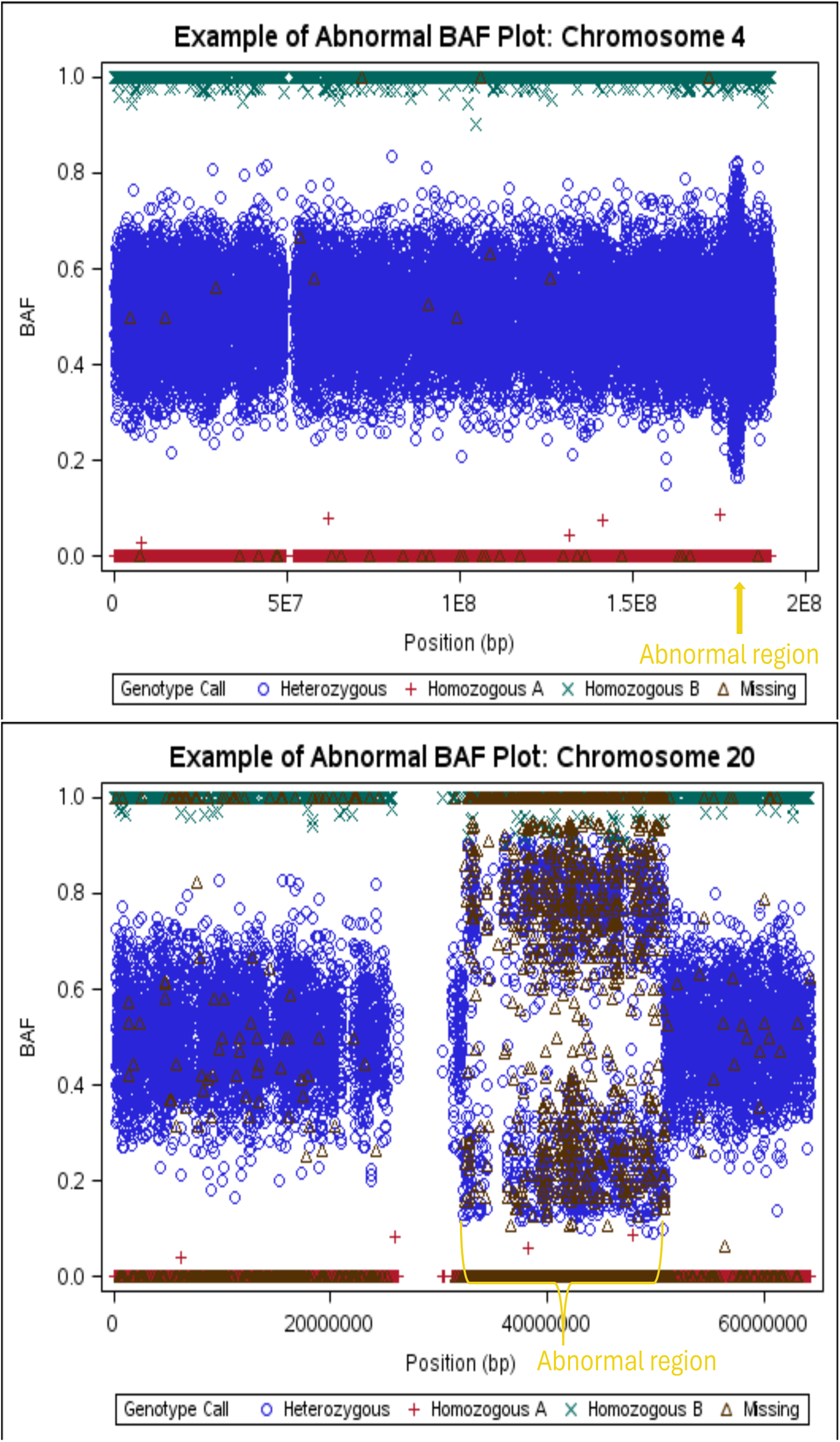
BAF for abnormal chromosomes kept (top) and dropped (bottom)

### Variant Mendel Error Filtering

After the B-allele Filtering described above, we checked for autosomal Mendel inconsistencies with the Loki Program^14^. For the sake of efficiency, we broke the genome down into sub-chunks with a few thousand markers each and ran them separately. We examined Mendel errors (ME) per marker and defined “pedcount” as the number of pedigrees in which a marker has any MEs (so a family with multiple MEs only adds 1 to the pedcount of a variant).

Since the goal of our study is to identify inherited variants, for the “clean” analysis data set, we filtered out “bad” markers as determined by a high pedcount. While a small number of MEs may represent de novo mutation, it is far more likely that they come from genotyping error: the de novo base pair mutation rate of 1.2E-8 is much smaller than expected genotype call error rate^15^. In a cancer study, one may make a different choice, but to identify germline genetic variants, this seemed like the better choice.

Variants were stratified by minor allele frequency (MAF): the ability to detect MEs depends on MAF, so each MAF strata had a different pedcount threshold. The threshold was set to remove the 0.5% of markers with the largest pedcount scores in each stratum. We have found this threshold to meet a utilitarian balance between eliminating problematic/artifactual variant calls and maintaining data on as many markers as possible. Table 4a contains the number of diallelic SNPs removed from each chromosome. Table 4b contains the number of diallelic INDELs removed from each chromosome, including the INDELs that overlap by at least one base pair position. The overlapping diallelic INDELs may not truly be diallelic and a multi-allelic QC is called for.

**Table 4a.** The number of sub-chunks per chunk per chromosome, number of SNPs before Mendel error QC, number of SNPs dropped, and number of SNPs remained after QC.

| Chr | # Of Chunks | # Of Sub-chunks | # SNPs before ME QC | # Monomorphic SNPs Dropped | # SNPs Dropped (pedcount & MAF > cutoff) | # SNPs Remained after QC | # SNPs Mendelian Inconsistency Zeroed-Out a Whole Pedigree |
| --- | --- | --- | --- | --- | --- | --- | --- |
| 1 | 20 | 668 | 4,618,096 | 25,853 | 14,635 | 4,577,608 | 33,079 |
| 2 | 20 | 724 | 4,993,363 | 28,913 | 11,888 | 4,952,562 | 24,776 |
| 3 | 16 | 603 | 4,160,996 | 22,078 | 10,627 | 4,128,291 | 23,199 |
| 4 | 16 | 581 | 4,017,409 | 28,366 | 7,955 | 3,981,088 | 17,024 |
| 5 | 15 | 538 | 3,724,815 | 19,702 | 7,627 | 3,697,486 | 15,480 |
| 6 | 14 | 516 | 3,563,155 | 20,580 | 7,780 | 3,534,795 | 16,011 |
| 7 | 13 | 484 | 3,354,431 | 21,635 | 9,281 | 3,323,515 | 19,884 |
| 8 | 12 | 465 | 3,215,525 | 16,145 | 5,995 | 3,193,385 | 11,855 |
| 9 | 12 | 371 | 2,557,569 | 16,102 | 7,433 | 2,534,034 | 17,476 |
| 10 | 11 | 411 | 2,837,527 | 13,410 | 7,566 | 2,816,551 | 16,226 |
| 11 | 11 | 412 | 2,841,624 | 17,184 | 6,427 | 2,818,013 | 13,228 |
| 12 | 11 | 397 | 2,734,575 | 15,581 | 6,759 | 2,712,235 | 12,869 |
| 13 | 9* | 292 | 2,011,463 | 11,140 | 4,763 | 1,995,560 | 10,526 |
| 14 | 8* | 268 | 1,847,584 | 11,203 | 4,109 | 1,832,272 | 8,582 |
| 15 | 8* | 246 | 1,692,150 | 10,399 | 4,393 | 1,677,358 | 9,295 |
| 16 | 8 | 280 | 1,924,107 | 10,086 | 5,569 | 1,908,452 | 11,849 |
| 17 | 7 | 243 | 1,682,421 | 8,932 | 5,785 | 1,667,704 | 11,571 |
| 18 | 7 | 233 | 1,595,922 | 9,299 | 3,459 | 1,583,164 | 6,638 |
| 19 | 5 | 191 | 1,326,246 | 6,838 | 5,574 | 1,313,834 | 11,409 |
| 20 | 6 | 198 | 1,366,602 | 8,746 | 6,017 | 1,351,839 | 15,153 |
| 21 | 4 | 121 | 831,437 | 4,233 | 3317 | 823,887 | 9,468 |
| 22 | 5 | 124 | 861,777 | 4,816 | 4751 | 852,210 | 12,429 |
| Total | 238 | 8,366 | 57,758,794 | 331,241 | 151,710 | 57,275,843 | 328,027 |

**Table 4b.** The number of sub-chunks per chunk per chromosome, number of INDELs before Mendel error QC, number of INDELs dropped, and number of INDELs remained after QC.

| Chr | # Of Chunks | # Of Sub-chunks | # INDELs before ME QC | # Monomorphic INDELs Dropped | # INDELs Dropped (pedcount & MAF > cutoff) | # Overlapping INDELs Dropped | # INDELs Remained after QC | # INDELs Mendelian Inconsistency Zeroed-Out a Whole Pedigree |
| --- | --- | --- | --- | --- | --- | --- | --- | --- |
| 1 | 20 | 73 | 422,515 | 22,720 | 1,459 | 24,796 | 373,540 | 2,613 |
| 2 | 20 | 77 | 456,697 | 25,445 | 1,315 | 27,069 | 402,868 | 2,231 |
| 3 | 16 | 63 | 378,968 | 20,765 | 982 | 21,962 | 335,259 | 1,803 |
| 4 | 16 | 60 | 370,598 | 20,115 | 927 | 21,426 | 328,130 | 1,519 |
| 5 | 15 | 56 | 338,387 | 18,803 | 808 | 19,566 | 299,210 | 1,333 |
| 6 | 14 | 54 | 336,509 | 17,623 | 926 | 19,722 | 298,238 | 1,854 |
| 7 | 13 | 50 | 304,805 | 15,653 | 1,037 | 17,541 | 270,574 | 2,044 |
| 8 | 12 | 46 | 273,601 | 15,172 | 724 | 16,248 | 241,457 | 1,042 |
| 9 | 12 | 38 | 221,325 | 11,911 | 742 | 13,115 | 195,557 | 1,600 |
| 10 | 11 | 42 | 253,666 | 13,158 | 897 | 15,698 | 223,913 | 1,643 |
| 11 | 11 | 42 | 250,859 | 13,654 | 765 | 14,670 | 221,770 | 1,345 |
| 12 | 11 | 42 | 255,125 | 13,299 | 716 | 15,093 | 226,017 | 1,332 |
| 13 | 9* | 32 | 193,134 | 10,709 | 521 | 11,095 | 170,809 | 989 |
| 14 | 8* | 29 | 170,696 | 8,997 | 451 | 9,828 | 151,420 | 836 |
| 15 | 8* | 27 | 155,814 | 8,648 | 402 | 8,898 | 137,866 | 906 |
| 16 | 8 | 25 | 151,153 | 7,475 | 563 | 9,044 | 134,071 | 1,138 |
| 17 | 7 | 26 | 156,946 | 6,947 | 720 | 9,352 | 139,927 | 1,284 |
| 18 | 7 | 25 | 148,224 | 8,705 | 416 | 9,033 | 130,070 | 673 |
| 19 | 5 | 19 | 121,428 | 4,236 | 562 | 6,615 | 110,015 | 1,188 |
| 20 | 6 | 21 | 119,541 | 6,516 | 583 | 7,481 | 104,961 | 1,300 |
| 21 | 4 | 13 | 74,920 | 3,402 | 392 | 4,825 | 66,301 | 872 |
| 22 | 5 | 13 | 74,642 | 3,253 | 525 | 4,656 | 66,208 | 1,205 |
| Total | 238 | 873 | 5,229,553 | 277,206 | 16,433 | 307,733 | 4,628,181 | 30,750 |

**\*\* There are 469,506 diallelic INDELs that overlap by at least one base pair position. 157,739 of these INDELs are also monomorphic and another 4,034 of these INDELs are on the list of variants with “pedcount & maf” > cutoff.**

Tables 5a and 5b show the distributions of variants by MAF strata and number of pedigrees with at least one ME. If a marker had a ME pedcount below the threshold for that marker’s MAF strata, we set that marker to missing in each family in which an error occurred, while retaining the data for that marker in families with no MEs.

**Table 5a.** Distribution of diallelic SNPs by MAF and number of pedigrees with at least one ME in that SNP (“pedcount”). Green highlight = no MEs, markers included. Yellow = Markers set to missing in families with MEs. Red = markers removed for excess MEs.

| Ped Count | MAF 0.4 to 0.5 | MAF 0.3 to 0.4 | MAF 0.2 to 0.3 | MAF 0.1 to 0.2 | MAF 0.05 to 0.1 | MAF 0.01 to 0.05 | MAF 0.005 to 0.01 | MAF 0.001 to 0.005 | MAF 0 to 0.001 |
| --- | --- | --- | --- | --- | --- | --- | --- | --- | --- |
| 0 | 865,518 | 927,675 | 1,063,086 | 1,420,629 | 1,127,394 | 2,432,204 | 1,274,865 | 5,760,825 | 42,075,620 |
| 1 | 11,018 | 10,795 | 9,830 | 8,691 | 4,488 | 11,878 | 16,603 | 101,119 | 98,057 |
| 2 | 760 | 733 | 763 | 852 | 658 | 9,087 | 15,779 | 38,796 | 5,774 |
| 3 | 346 | 263 | 336 | 342 | 414 | 10,783 | 12,448 | 12,236 | 329 |
| 4 | 139 | 149 | 190 | 180 | 260 | 11,448 | 7,448 | 3,254 | 23 |
| 5 | 108 | 96 | 110 | 115 | 229 | 11,045 | 3,981 | 794 | 1 |
| 6 | 71 | 59 | 80 | 85 | 239 | 10,069 | 1,597 | 204 |  |
| 7 | 52 | 52 | 63 | 67 | 239 | 8,715 | 641 | 53 |  |
| 8 | 38 | 42 | 48 | 36 | 257 | 7,329 | 205 | 17 |  |
| 9 | 32 | 28 | 26 | 36 | 254 | 5,936 | 81 | 11 |  |
| 10 | 22 | 17 | 20 | 29 | 272 | 4,722 | 28 | 4 |  |
| 11-50 | 184 | 146 | 106 | 124 | 1955 | 12617 | 33 | 8 |  |
| 51-100 | 10 | 4 | 8 | 2 | 7 | 4 |  |  |  |
| 101 | 1 |  |  |  |  |  |  |  |  |
| 106 |  |  | 1 |  |  |  |  |  |  |
| 116 |  |  |  |  | 1 |  |  |  |  |
| 137 |  | 1 |  |  |  |  |  |  |  |
| 150 |  |  |  | 1 |  |  |  |  |  |
| Total | 878,299 | 940,060 | 1,074,667 | 1,431,189 | 1,136,667 | 2,535,837 | 1,333,709 | 5,917,321 | 42,179,804 |

**Table 5b.**
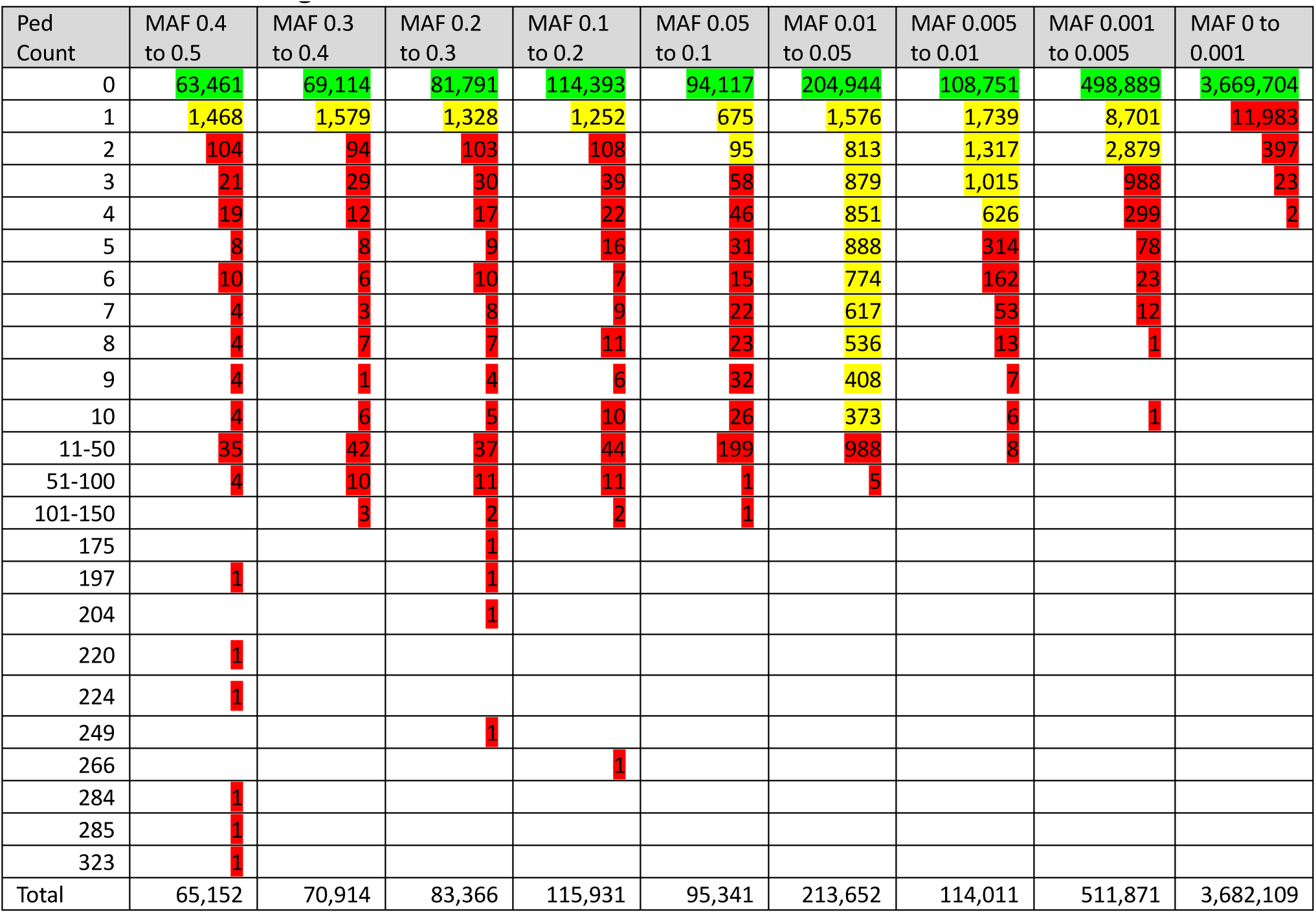
Distribution of diallelic INDELs by MAF and number of pedigrees with at least one ME in that INDEL (“pedcount”). Green highlight = no MEs, markers included. Yellow = Markers set to missing in families with MEs. Red = markers removed for excess MEs.

| Ped Count | MAF 0.4 to 0.5 | MAF 0.3 to 0.4 | MAF 0.2 to 0.3 | MAF 0.1 to 0.2 | MAF 0.05 to 0.1 | MAF 0.01 to 0.05 | MAF 0.005 to 0.01 | MAF 0.001 to 0.005 | MAF 0 to 0.001 |
| --- | --- | --- | --- | --- | --- | --- | --- | --- | --- |
| 0 | 63,461 | 69,114 | 81,791 | 114,393 | 94,117 | 204,944 | 108,751 | 498,889 | 3,669,704 |
| 1 | 1,468 | 1,579 | 1,328 | 1,252 | 675 | 1,576 | 1,739 | 8,701 | 11,983 |
| 2 | 104 | 94 | 103 | 108 | 95 | 813 | 1,317 | 2,879 | 397 |
| 3 | 21 | 29 | 30 | 39 | 58 | 879 | 1,015 | 988 | 23 |
| 4 | 19 | 12 | 17 | 22 | 46 | 851 | 626 | 299 | 2 |
| 5 | 8 | 8 | 9 | 16 | 31 | 888 | 314 | 78 |  |
| 6 | 10 | 6 | 10 | 7 | 15 | 774 | 162 | 23 |  |
| 7 | 4 | 3 | 8 | 9 | 22 | 617 | 53 | 12 |  |
| 8 | 4 | 7 | 7 | 11 | 23 | 536 | 13 | 1 |  |
| 9 | 4 | 1 | 4 | 6 | 32 | 408 | 7 |  |  |
| 10 | 4 | 6 | 5 | 10 | 26 | 373 | 6 | 1 |  |
| 11-50 | 35 | 42 | 37 | 44 | 199 | 988 | 8 |  |  |
| 51-100 | 4 | 10 | 11 | 11 | 1 | 5 |  |  |  |
| 101-150 |  | 3 | 2 | 2 | 1 |  |  |  |  |
| 175 |  |  | 1 |  |  |  |  |  |  |
| 197 | 1 |  | 1 |  |  |  |  |  |  |
| 204 |  |  | 1 |  |  |  |  |  |  |
| 220 | 1 |  |  |  |  |  |  |  |  |
| 224 | 1 |  |  |  |  |  |  |  |  |
| 249 |  |  | 1 |  |  |  |  |  |  |
| 266 |  |  |  | 1 |  |  |  |  |  |
| 284 | 1 |  |  |  |  |  |  |  |  |
| 285 | 1 |  |  |  |  |  |  |  |  |
| 323 | 1 |  |  |  |  |  |  |  |  |
| Total | 65,152 | 70,914 | 83,366 | 115,931 | 95,341 | 213,652 | 114,011 | 511,871 | 3,682,109 |

### Mitochondrial (Mt) chromosome filtering

The processing of the mitochondrial chromosome is perhaps the most different from the autosomal chromosomes. Not only is it matrilineal (for both male and female subjects), but because there are many copies of it per cell, the depth of coverage is vastly greater: in the thousands for most sites (figure 4). As a result, no sites were removed due to low coverage. Less than 1% of calls have relatively low depth (<100), all from the D-loop region; use variants from this region with caution. As we noted above, the Mt variants were called as haploid.

**Figure 4.**
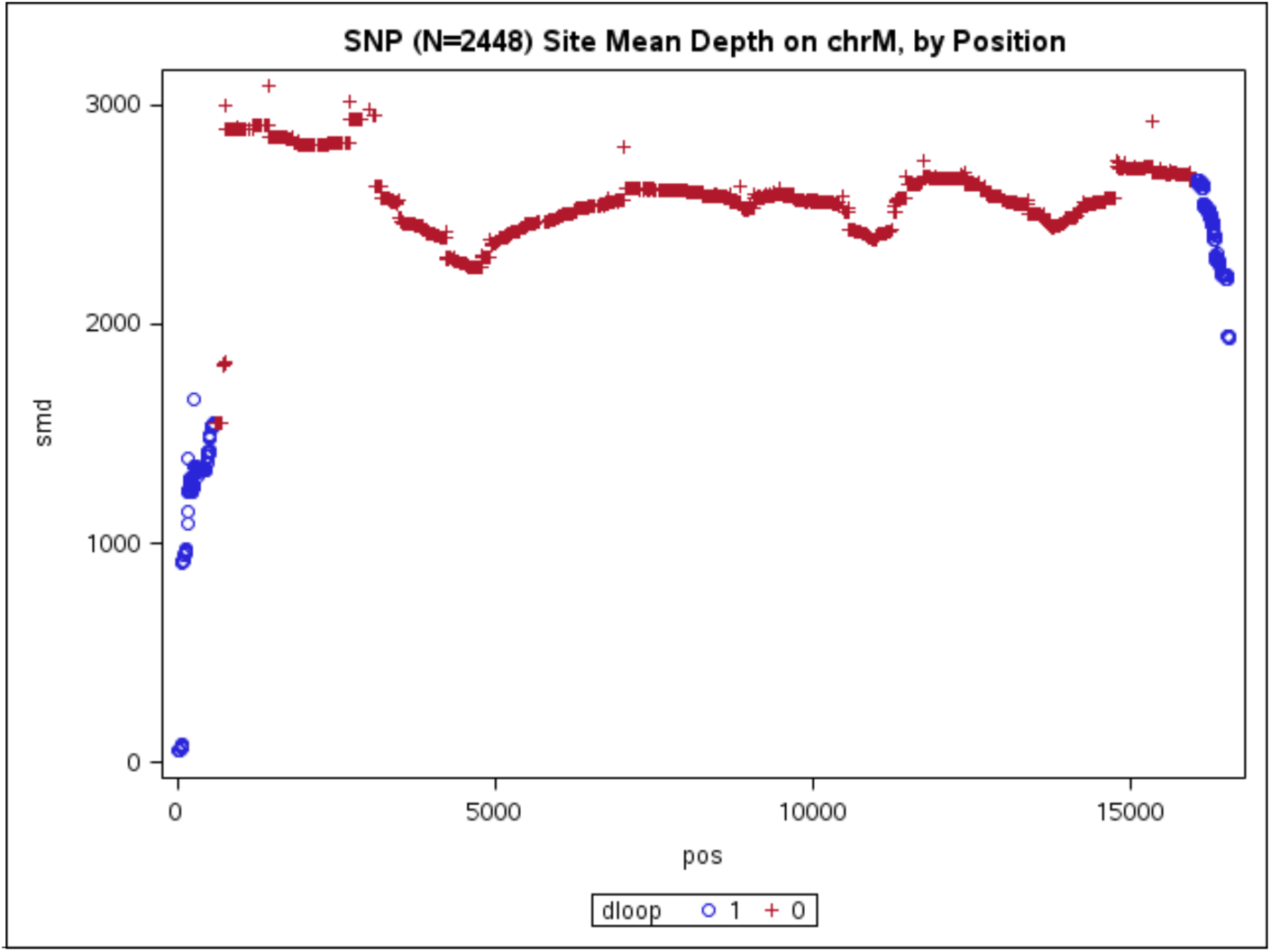
Chromosome Mt Site Mean Depth

Most variants on Mt are rare; in LLFS, 27% of SNPs and 50% of INDELs are singletons. Mt contains only 63 common (MAF>=5%) SNPs. There is very little missingness in the Mt calls (0.0002%), and this occurs in the variants in the D-loop region.

As one has Mendel errors in autosomal variants, one can check for lineage errors via the matrilineal line for Mt variants. A lineage can only be checked if it contains at least two members for comparison. Only 2,642 subjects were in a ‘checkable’ matrilineal line (598 unique lines). 101 SNPs (out of 2,439) and 2 INDELS (out of 18) had at least one Mt lineage error (Table 6).

**Table 6.** Number of SNPs/INDELs with Mt Lineage Errors.

| Number of SNPs | # of lineage errors | Number of INDELs | # of lineage errors |
| --- | --- | --- | --- |
| 2,338 | 0 | 16 | 0 |
| 91 | 1 | 1 | 1 |
| 4 | 2 | 1 | 5 |
| 4 | 3 |  |  |
| 1 | 4 |  |  |
| 1 | 35 |  |  |

Most lineage errors occurred in the D-loop region. One SNP and one INDEL (both listed above) were removed due to lineage errors. For the remaining lineage errors, we set each variant to missing in the lineage in which the error occurred, while retaining the data for that variant in lineages with no errors.

### Y chromosome filtering

As noted above, the regions exclusive to the Y chromosome (chrY) were called as haploid (similar to Mt) and only in male subjects by using GATK. The regions shared with X (PAR1, PAR2, and XTR) were handled in the processing of the X chromosome (see below). Note that the typical depth of coverage for the Y chromosome was roughly half that of the autosomes since most males have one copy of their Y chromosome compared to two copies of each autosome. Consequently, individual Y variant calls were filtered out if depth was less than 10 (at least ten sequence reads covering the variant site) or greater than 150. This filter was selected to balance the competing needs of quality data and complete data. Either low or high depth can be an indicator that the variant could be an artifact. In particular, there are spikes of extreme depth in known “blacklisted” regions around the centromere (∼10.4Mbp) and the q-arm of chrY contains many ampliconic and palindromic sequences making it difficult to sequence and resulting in large regions of missing data, as seen in Figure 7. Calls that do not meet these basic filter criteria were set to missing.

**Figure 5.**
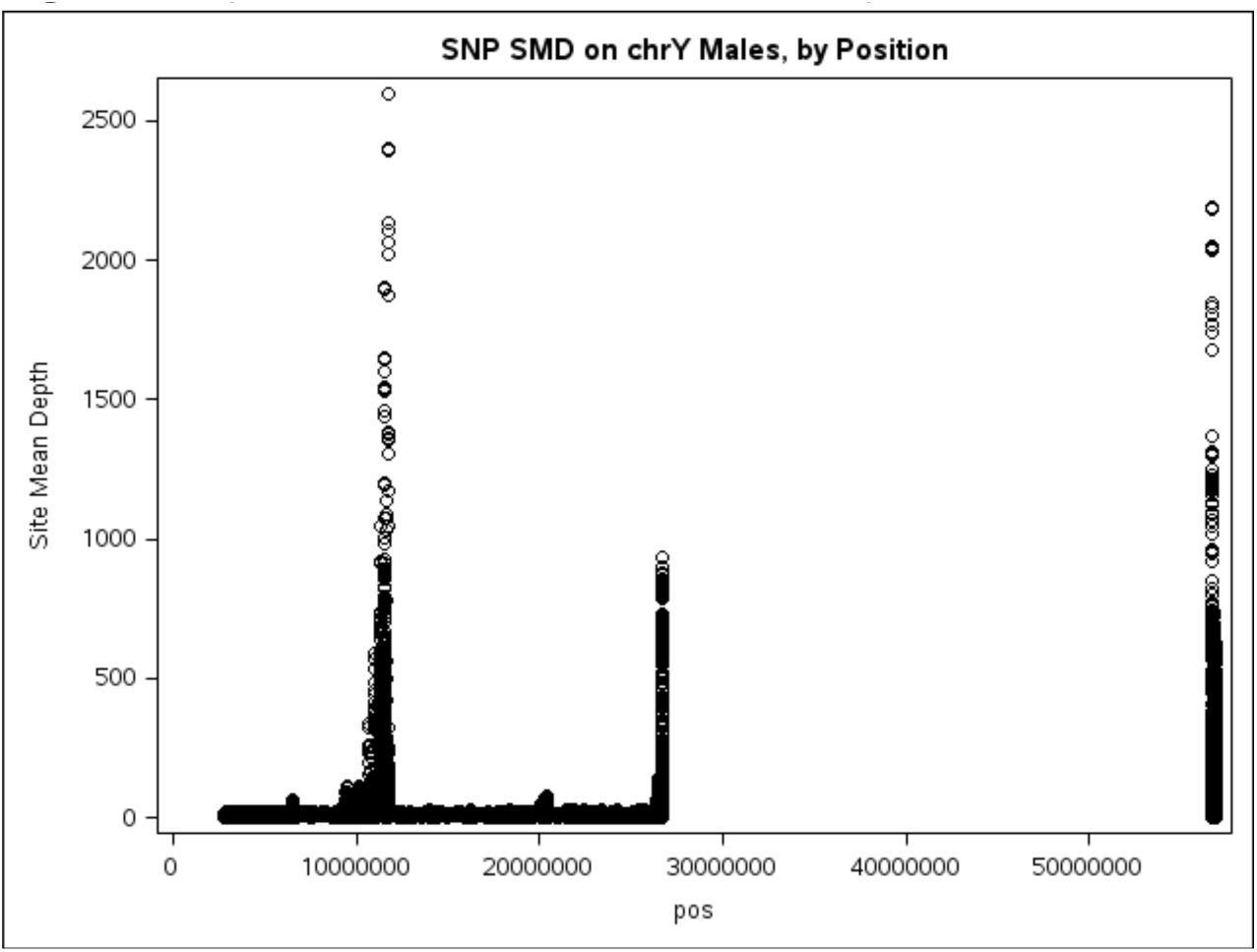
Spikes of Extreme Site Mean Depth on Chromosome Y

As with the autosomes, due to the removal of subjects, and setting variant calls to missing based on the depth filters, there were several variants that became monomorphic or completely missing. We removed these non-informative variants.

### Low call rate on Y

Low call rate can be indicative of poor sample quality or other issues. In particular for chrY, low call rate may also be due to loss of the Y chromosome in older subjects. In LLFS, older males have a higher missing rate than younger males, an effect that we did not want to factor into our call rate filter. Therefore, to avoid bias, we selected a ‘base’ set of 1,092 male subjects (age<70 and call rate>0.8; see Figure 6) and used this set of subjects to calculate the per-variant call rate. This call rate statistic should be unaffected by loss of Y in older males.

**Figure 6.**
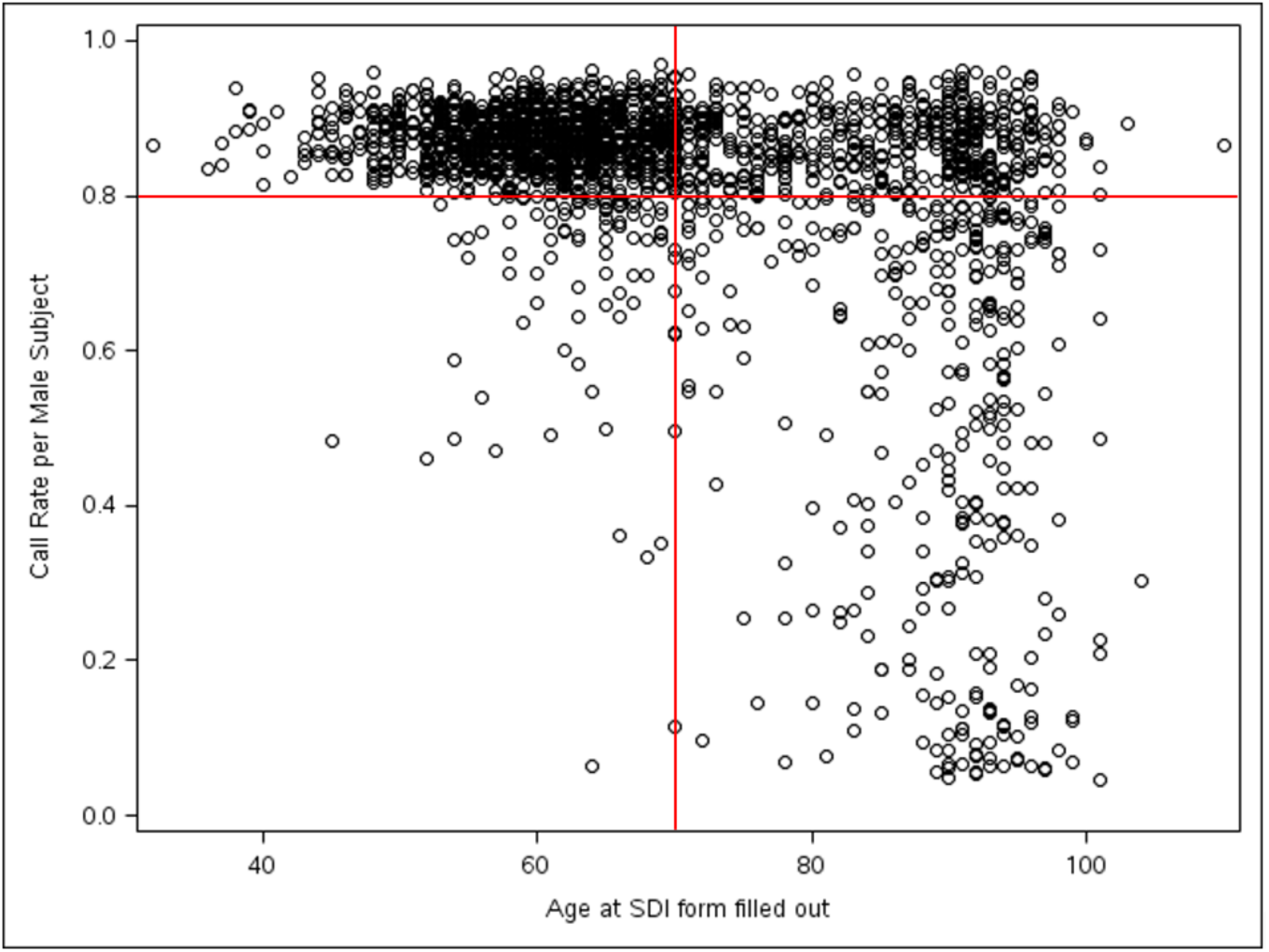
“Base” Subjects for calculating per-variant call rate on Chromosome Y: only subjects in upper left used to establish per-variant call rates for filtering

After reviewing histograms and summary statistics, we filtered variants with a call rate of less than 80%. This allowed us to remove extreme outliers, while still retaining approximately 90% of the variants. 33.7% of the variants removed by this call rate filter are in “blacklisted” regions, compared to only 11.6% of the remaining variants. Table 7 shows the number of variants removed during each QC step.

**Table 7.** Preliminary QC Stats on Diallelic SNPs and INDELs in Chromosome Y.

| Chr | Type | # Variants before QC | % calls set to missing due to depth filters | # (Additional) Variants Dropped |  |  |  | After Preliminary QC |
| --- | --- | --- | --- | --- | --- | --- | --- | --- |
|  |  |  |  | All Missing or Monomorphic |  | “Base Sub” Call Rate <0.8 |  |  |
|  |  |  |  | N | % | N | % |  |
| Y | SNP | 176,085 | 21.0 | 85,155 | 48.4 | 9,679 | 10.6 | 81,251 |
| Y | INDEL | 23,224 | 25.0 | 14,064 | 60.6 | 1,435 | 15.7 | 7,725 |

### Chromosome Y Lineage Errors

We checked for lineage errors via the patrilineal line (males only), if available. Only 971 male subjects were in a ‘checkable’ patrilineal line (307 unique lines). 9,519 SNPs (out of 81,251) and 762 INDELs (out of 7,725) had at least one chrY lineage error.

We used a similar method to remove chrY lineage errors to that we used to remove MEs in the autosome (above). Markers with the most lineage errors were completely removed, while markers with fewer lineage errors were set to missing in specific lineages in which the error occurred, while retaining the data for that marker in lineages with no errors. Tables 8a and 8b show the distributions of variants by MAF and number of pedigrees with at least one lineage error.

**Table 8a.** Distribution of diallelic chromosome Y SNPs by MAF and number of pedigrees with at least one lineage error (LE) in that SNP (“pedcount”). Green highlight = no LEs, markers included. Yellow = Markers set to missing in families with LEs. Red = markers removed for excess LEs.

| Ped Count | MAF 0.4 to 0.5 | MAF 0.3 to 0.4 | MAF 0.2 to 0.3 | MAF 0.1 to 0.2 | MAF 0.05 to 0.1 | MAF 0.01 to 0.05 | MAF 0.005 to 0.01 | MAF 0.001 to 0.005 | MAF 0 to 0.001 |
| --- | --- | --- | --- | --- | --- | --- | --- | --- | --- |
| 0 | 576 | 79 | 195 | 510 | 1,712 | 3,747 | 2,993 | 25,390 | 36,530 |
| 1 | 5 | 1 | 1 | 3 | 13 | 35 | 42 | 847 | 1,713 |
| 2 |  | 2 |  | 4 | 8 | 24 | 40 | 580 | 72 |
| 3 | 1 |  |  | 1 | 4 | 19 | 67 | 388 |  |
| 4 | 1 |  |  | 2 | 3 | 21 | 84 | 224 |  |
| 5 |  |  | 1 |  | 2 | 29 | 122 | 136 |  |
| 6 |  |  |  |  |  | 35 | 98 | 60 |  |
| 7 |  |  |  | 1 | 4 | 30 | 122 | 29 |  |
| 8 |  |  | 1 | 2 |  | 69 | 112 | 7 |  |
| 9 |  |  | 1 | 1 |  | 65 | 80 | 2 |  |
| 10 |  |  |  |  |  | 60 | 52 |  |  |
| 11-100 | 16 | 16 | 45 | 449 | 698 | 1,361 | 74 |  |  |
| 101-200 | 379 | 478 | 435 | 231 |  |  |  |  |  |
| 201 | 2 |  |  |  |  |  |  |  |  |
| 202 | 1 |  |  |  |  |  |  |  |  |
| 203 | 3 |  |  |  |  |  |  |  |  |
| 204 | 1 |  |  |  |  |  |  |  |  |
| 205 | 2 |  |  |  |  |  |  |  |  |
| 208 | 2 |  |  |  |  |  |  |  |  |
| Total | 989 | 576 | 679 | 1,204 | 2,444 | 5,495 | 3,886 | 27,663 | 38,315 |

**Table 8b.** Distribution of diallelic chromosome Y INDELs by MAF and number of pedigrees with at least one lineage error (LE) in that INDEL (“pedcount”). Green highlight = no LEs, markers included. Yellow = Markers set to missing in families with LEs. Red = markers removed for excess LEs.

| Ped Count | MAF 0.4 to 0.5 | MAF 0.3 to 0.4 | MAF 0.2 to 0.3 | MAF 0.1 to 0.2 | MAF 0.05 to 0.1 | MAF 0.01 to 0.05 | MAF 0.005 to 0.01 | MAF 0.001 to 0.005 | MAF 0 to 0.001 |
| --- | --- | --- | --- | --- | --- | --- | --- | --- | --- |
| 0 | 45 | 4 | 15 | 59 | 165 | 400 | 336 | 2,627 | 3,312 |
| 1 |  |  |  |  | 1 | 7 | 10 | 86 | 139 |
| 2 |  |  |  |  |  | 1 | 7 | 56 | 6 |
| 3 |  |  |  |  | 2 | 2 | 7 | 34 |  |
| 4 |  |  |  |  |  | 5 | 8 | 15 |  |
| 5 |  |  |  |  |  |  | 9 | 10 |  |
| 6 |  |  |  |  |  |  | 9 | 2 |  |
| 7 |  |  |  |  |  | 2 | 8 |  |  |
| 8 |  |  |  |  |  |  | 12 | 2 |  |
| 9 |  |  |  |  |  | 4 | 5 |  |  |
| 10 |  |  |  |  |  | 7 | 4 |  |  |
| 11-100 |  |  | 3 | 39 | 36 | 84 | 3 |  |  |
| 101-200 | 30 | 37 | 46 | 21 |  |  |  |  |  |
| 203 | 1 |  |  |  |  |  |  |  |  |
| 204 | 1 |  |  |  |  |  |  |  |  |
| 205 |  | 1 |  |  |  |  |  |  |  |
| Total | 77 | 42 | 64 | 119 | 204 | 512 | 418 | 2,832 | 3,457 |

### Chromosome X processing

For Chromosome X (chrX), we used a mix of diploid and haploid calling. Female subjects were called as diploid, along with males for variants in the pseudo-autosomal regions (PAR). Male subjects in non-PAR regions on chromosome X were called as haploid. It appeared that sequence reads were not mapped to the X-transposed region (XTR) on chrY, so male XTR variants are not included in this release. There did not appear to be an issue with XTR sequence on chrX, so female XTR calls are included.

Diallelic SNPs and INDELs on chromosome X were processed separately, as with the autosomes. The raw calls with our “mixed” haploid/diploid application of GATK produced 2,762,116 SNPs and 274,110 INDELs on chrX. SNPs in “blacklisted” regions account for 31% and 34% of chrX SNP and INDEL calls, respectively. Table 9 shows the number of variants in each region.

**Table 9.** Number of Chromosome X Variants, by Region and Sex.

| Region | Female | Male | N SNPs | N INDELs |
| --- | --- | --- | --- | --- |
| Pure X | Diploid | Haploid | 2,576,816 | 253,484 |
| PAR1 | Diploid | Diploid | 91,514 | 11,724 |
| XTR | Diploid | NA (not released) | 84,906 | 8,012 |
| PAR2 | Diploid | Diploid | 8,880 | 890 |

Depth statistics varied on chrX, depending on sex, ploidy, and region. In particular, there are spikes of extreme depth in known “blacklisted” region, as seen in Figure 7. For females (entire chromosome) and PAR regions for males, for each sequence call at each variant site, the individual call was filtered out for depth less than 20 or greater than 300, the same as we used for the autosomes. For males in non-PAR regions, the individual call was filtered out for depth less than 10 or greater than 150, as we did for regions unique to chrY.

These filters were selected to balance the competing needs of quality data and complete data. With low depth, there is high risk that both haplotypes are not represented or that the variant is an artifact, while with high depth, there is high risk that the variant is an artifact. Calls that do not meet these basic filter criteria were set to missing.

**Figure 7.**
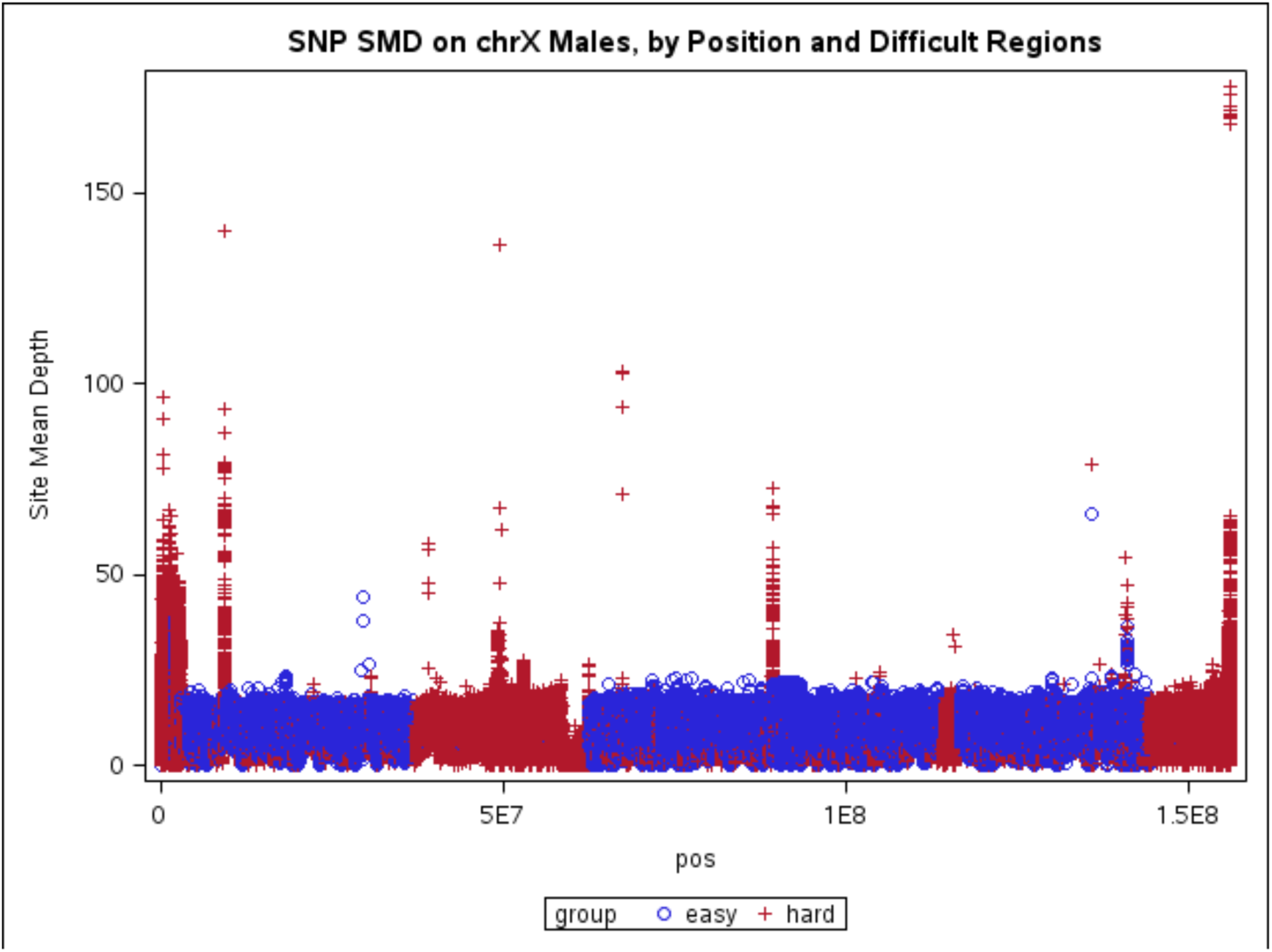
Spikes of Extreme Site Mean Depth on Chromosome X

Due to filtering, some variants that became either monomorphic or completely missing. As on other chromosomes, we removed these non-informative variants.

### Excess heterozygosity

We examined heterozygosity statistics separately for males and females, and by region (e.g., for males, heterozygosity only exists in PAR regions on chrX); see above for general discussion of heterozygosity filters. Figures 8a and 8b show SNP heterozygosity by HWE p-value for females, before and after the excess heterozygosity filter. INDEL heterozygosity figures look similar and are not shown here.

**Figures 8a-8b.**
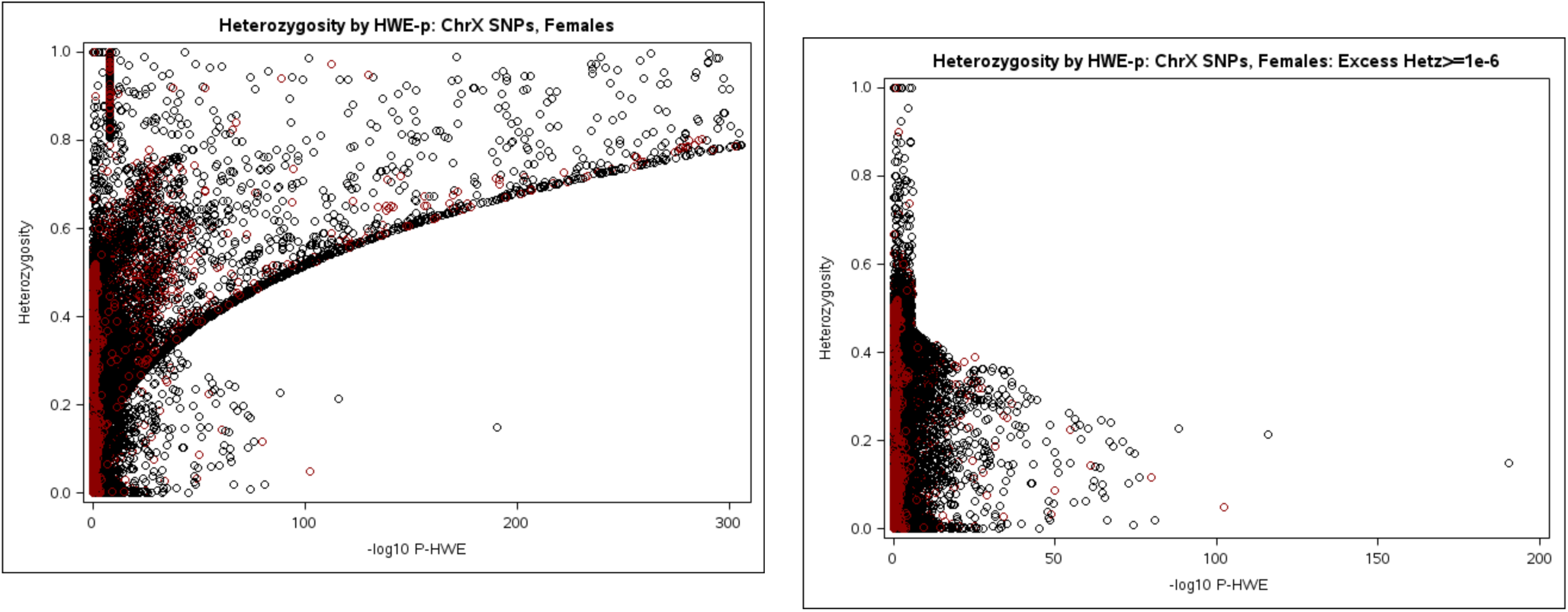
Chromosome X Before and After Excess Heterozygosity Filter

In the non-PAR regions (where heterozygosity only exists for females), we chose to filter out variants which failed the HWE test because of excess heterozygosity with p-value <1e-6. Heterozygosity statistics were calculated from females only, but removed for all subjects. There were still some markers with heterozygosity > 0.55, and we subsequently removed those as well.

In the PAR regions (where heterozygosity exists for males and females), we calculated heterozygosity in males, females, and the combined sample, and then removed variants if excess heterozygosity <1E-6 or heterozygosity>0.55 in any of those three groups. Table 10 shows the number of variants removed on chrX due to heterozygosity filters.

**Table 10.**
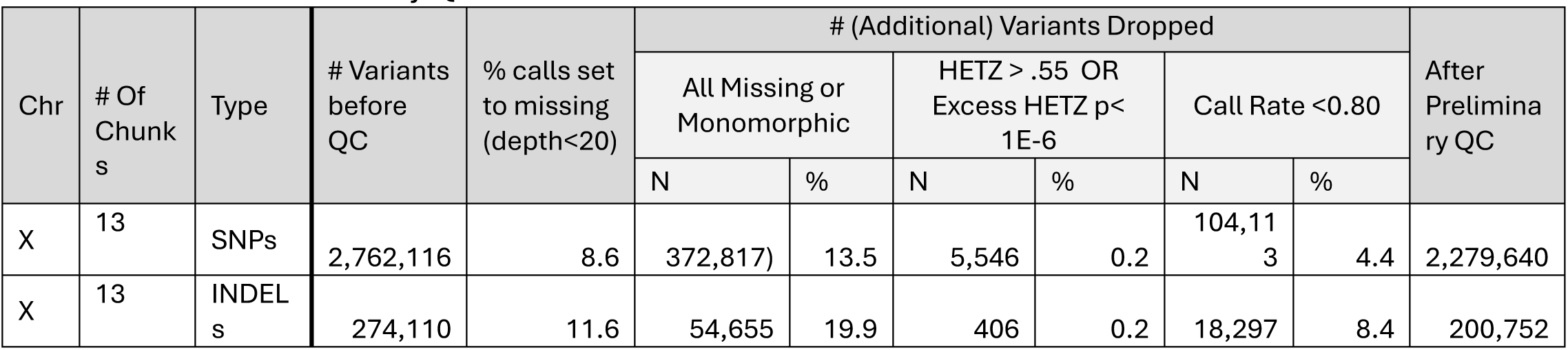
Preliminary QC Stats on Diallelic SNPs and INDELs in Chromosome X VCF Files.

| Chr | # Of<br>Chunk<br>s | Type | # Variants<br>before<br>QC | % calls set<br>to missing<br>(depth<20) | # (Additional) Variants Dropped |  |  |  |  |  | After<br>Prelimina<br>ry QC |
| --- | --- | --- | --- | --- | --- | --- | --- | --- | --- | --- | --- |
|  |  |  |  |  | All Missing or<br>Monomorphic |  | HETZ > .55 OR<br>Excess HETZ p<<br>1E-6 |  | Call Rate <0.80 |  |  |
|  |  |  |  |  | N | % | N | % | N | % |  |
| X | 13 | SNPs | 2,762,116 | 8.6 | 372,817) | 13.5 | 5,546 | 0.2 | 104,113 | 4.4 | 2,279,640 |
| X | 13 | INDEL<br>s | 274,110 | 11.6 | 54,655 | 19.9 | 406 | 0.2 | 18,297 | 8.4 | 200,752 |

As with the autosomal variants showing excess homozygosity, we did not remove such variants on chrX, since they could be real variants. However, they could be artifacts and even if they are real, standard association tests are known to yield biased results with such variants. Thus, care should be exercised in using variants with excess homozygosity.

### Variant Call Rate

Low call rate can be indicative of poor sample quality or other issues. Variants with many lower depth calls now have higher missing rates, since calls with low depth were set to missing. Call rate was calculated in males, females, and the combined sample. After reviewing histograms and summary statistics, we filtered variants with a call rate of less than 80% in any of those 3 groups. This allowed us to remove extreme outliers, while still retaining approximately 95% of the variants.

### Chromosome X Mendel Errors

We identified chrX MEs using PLINK^16^ (options –mendel-duos and –split-x); 28,717 SNPs (out of 2,279,640) and 2,650 INDELs (out of 200,752) had at least one chrX ME.

We used the same method to remove chrX MEs that we used in the autosome. In short, markers with the most MEs were completely removed from the CSV files, while markers with fewer MEs were set to missing in specific families in which an error occurred, while retaining the data for that marker in families with no errors. Tables 11a and 11b show the distributions of variants by MAF and number of pedigrees with at least one ME.

**Table 11a.** Distribution of diallelic chromosome X SNPs by MAF and number of pedigrees with at least one ME in that SNP (“pedcount”). Green highlight = no MEs, markers included. Yellow = Markers set to missing in families with MEs. Red = markers removed for excess MEs.

| Ped Count | MAF 0.4 to 0.5 | MAF 0.3 to 0.4 | MAF 0.2 to 0.3 | MAF 0.1 to 0.2 | MAF 0.05 to 0.1 | MAF 0.01 to 0.05 | MAF 0.005 to 0.01 | MAF 0.001 to 0.005 | MAF 0 to 0.001 |
| --- | --- | --- | --- | --- | --- | --- | --- | --- | --- |
| 0 | 25,279 | 28,371 | 34,087 | 41,842 | 33,900 | 86,846 | 54,102 | 295,148 | 1,651,348 |
| 1 | 1,364 | 1,405 | 1,461 | 1,320 | 512 | 751 | 989 | 5,830 | 5,529 |
| 2 | 322 | 299 | 284 | 300 | 109 | 475 | 801 | 1,581 | 472 |
| 3 | 57 | 63 | 65 | 59 | 24 | 484 | 444 | 390 | 102 |
| 4 | 27 | 29 | 36 | 28 | 15 | 461 | 231 | 94 | 63 |
| 5 | 5 | 7 | 13 | 9 | 9 | 406 | 87 | 24 | 2 |
| 6 | 7 | 3 | 1 | 7 | 14 | 338 | 30 | 7 |  |
| 7 | 4 | 4 | 1 | 9 | 6 | 264 | 6 | 2 |  |
| 8 |  | 4 | 3 |  | 7 | 213 | 1 | 2 |  |
| 9 | 1 | 3 | 1 | 4 | 8 | 170 | 1 |  |  |
| 10 | 2 | 1 | 2 | 1 | 7 | 109 |  |  |  |
| 11-50 | 7 | 17 | 19 | 6 | 46 | 322 | 7 |  |  |
| 51-100 | 1 |  |  |  |  | 7 |  |  |  |
| 104 |  |  |  |  |  | 1 |  |  |  |
| 105 |  |  |  |  |  | 1 |  |  |  |
| 134 |  |  |  |  |  | 1 |  |  |  |
| 168 |  |  |  |  | 1 |  |  |  |  |
| Total | 27,076 | 30,206 | 35,973 | 43,585 | 34,658 | 90,849 | 56,699 | 303,078 | 1,657,516 |

**Table 11b.**
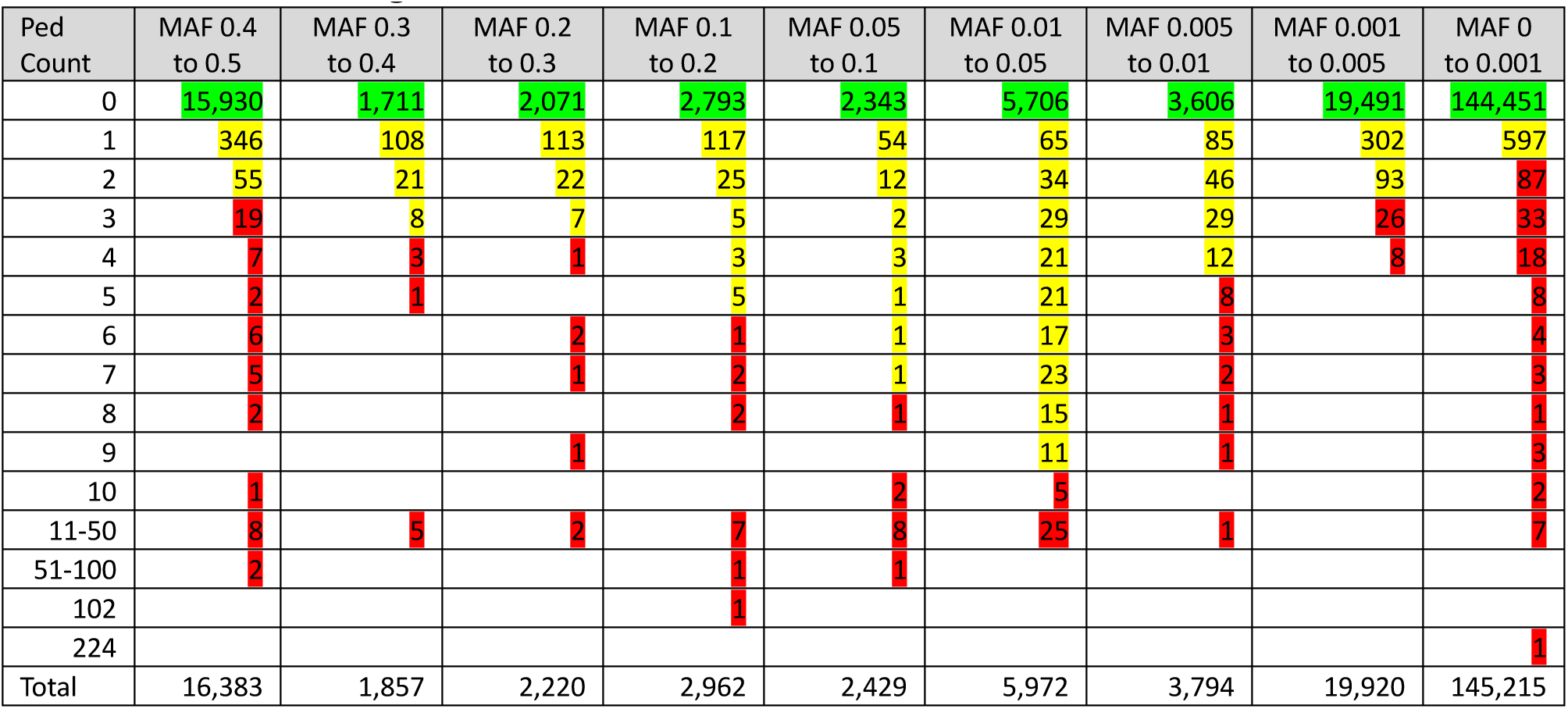
Distribution of diallelic chromosome X INDELs by MAF and number of pedigrees with at least one ME in that INDEL (“pedcount”). Green highlight = no MEs, markers. Yellow = Markers set to missing in families with MEs. Red = markers removed for excess MEs.

### Autosomal Linkage Markers and Identity by Descent Calculation

As LLFS is a family study, we are naturally interested in conducting linkage analysis. Linkage in families is a powerful tool: It is possible to identify causal genes with ∼100 (or sometimes even fewer) individuals in large families. However, linkage methods were developed in an era when genetic markers were sparser in the genome. The Lander-Green algorithm is linear in the number of genetic markers and running it with thousands of markers is practical. However, running with millions of sequence SNPs takes a thousand times longer and is not practical. Furthermore, in going from thousands of SNPs to millions, much of the information from the additional SNPs is essentially redundant due to chromosomes being inherited in blocks from each parent. This means that using every diallelic sequence SNP in a linkage analysis would vastly increase the computational burden while only providing a tiny increase in linkage information over a carefully selected set of thousands of SNPs. Thus, there is a clear computational advantage for using a carefully selected subset of sequence SNPs for linkage. We have focused primarily on autosomal linkage as tools for X chromosome linkage are more limited.

In addition to WGS data, we have GWAS chip data for most subjects. This includes a small number of family members for whom we have SNP chip, but not sequence data. Pedigree members with either GWAS chip or Sequence data can contribute to a linkage analysis, and we want to use both to maximize the information available for analysis. Having decided to only use a subset of SNPs, we decided to use the SNPs common to both our WGS and GWAS chip datasets as a starting point for choosing our linkage subset. We used only SNPs where we could match the alleles (sometimes after flipping) and frequencies, which resulted in a list of 517,020 SNPs. Most subjects have both GWAS chip and sequence data, so we computed kappa coefficients (a standard method for evaluating test/re-test data) for these markers. Most (99.75% or 515,764) SNPs had kappa > 0.95, and we decided to only use SNPs with kappa >0.999 (see table 12). For the very small number of calls where the GWAS and WGS calls differed, we used the GWAS call.

**Table 12.** Kappa Coefficient (> 0.95) GWAS vs. WGS.

| Category | Frequency | Percent | Cumulative Frequency |
| --- | --- | --- | --- |
| 0.95 < Kappa <= 0.96 | 146 | 0.03 | 146 |
| 0.96 < Kappa <= 0.97 | 261 | 0.05 | 407 |
| 0.97 < Kappa <= 0.98 | 468 | 0.09 | 875 |
| 0.98 < Kappa <= 0.99 | 1063 | 0.21 | 1938 |
| 0.99 < Kappa <= 0.999 | 17152 | 3.33 | 19090 |
| 0.999 < Kappa <= 1 | 496674 | 96.3 | 515764 |

While we had checked both the GWAS and the WGS data for Mendel errors, combining the data had the potential to introduce new errors, so we re-checked the combined data set with Loki and removed any errors.

To obtain centiMorgan (cM) positions for all these SNPs, we used the deCODE sex-specific SNP meiotic map^17^. If a SNP was in the map, we used the map position. If a SNP was not in the deCODE meiotic map, we used linear interpolation with bp position on SNPs that were in the map. There were a small number of SNPs where the bp order in GRCh38 did not agree with the cM position order in the deCODE map (due to being based on a different build); these SNPs were dropped.

To be useful for linkage, a marker must have some within family variation. We computed the average of the within family variation for each SNP and selected those with a value >0.1.

One problem with individual SNPs is that they are not very informative as linkage markers. Classical microsatellite markers had heterozygosities of ∼0.8, but the maximum for a diallelic SNP is 0.5 and most are much smaller. While one can use more SNPs to compensate, this comes at a computational burden. We decided to instead construct SNP haplotypes of five tightly linked SNPs and use these to span the genome. These constructed haplotypes have heterozygosity similar to those of microsatellites.

To select SNPs for the haplotype construction, we divided the genome into 0.5cM bins. Most bins had at least 5 SNPs (some had hundreds), but a few had less than five. If there were five or more SNPs in a bin, we used the first five in the bin. If there were less, we used only the SNPs in the bin, resulting in a small number of bins with haplotypes of fewer than five SNPs.

To generate the haplotypes in each bin, we used the ZAPLO program^18^, which uses pedigrees to construct haplotypes under the assumption of zero recombination. The SNPs in each bin will be less than 0.5cM apart, and sometimes much less than that, particularly in bins with hundreds of SNPs. However, a recombination is not impossible, and some will occur in the thousands of samples looking across the whole genome in our study. As a result, ZAPLO is not always able to find a zero-recombination haplotype configuration for each family in each bin. In such cases, we skipped over any family in a bin that failed to find a configuration, i.e., the data for that family in that bin was left as missing. However, we wanted to have at least a 99% completion rate for the haplotyping, so if the family skipping resulted in a missing rate of 1% or more, we re-selected the SNPs in the bin, according to the number of SNPs in the bin as follows:

a. if number of available SNPs in a bin > 20, move over by 5 SNPs (select 5 SNPs with choice 6 - 10, then 11 - 15, etc.)
b. if 10 < number of available SNPs in a bin is > 10 and <= 20, move over by 1 SNP (select 5 SNPs with choice 2 - 6, then 3 - 7, 4 - 8, etc.)
c. if number of available SNPs in a bin is > 6 and <= 10, drop 1, add 1, then move (select 5 SNPs with choice 2 - 6, then (1, 3 - 6), (1, 2, 4 - 6), etc.)
d. if number of available SNPs in a bin <= 5, try sets of N-1 (then N-2, etc.)

Note that ZAPLO will impute haplotypes in subjects with no data. We did not want to use this imputed data, so we removed the imputed haplotypes for subjects with no GWAS or WGS data. We did, however, use the haplotypes for subjects who had genetic data but might have been missing a called genotype at one of the SNPs in the haplotype. For the 6586 bins of 0.5cM that spanned the whole autosome, the number of SNPs per haplotype is in table 13. Note that 26 bins had no SNPs, and so no haplotype was computed for these bins.

**Table 13.** Number of SNPs per Bin.

| Number of SNPs per BIN | Number of BINs |
| --- | --- |
| 0 | 26 |
| 1 | 15 |
| 2 | 17 |
| 3 | 24 |
| 4 | 26 |
| 5 | 6478 |
| total | 6586 |

These haplotype markers can be used with most linkage programs (e.g. SOLAR^19^, MERLIN^20^, or Loki^14^) directly and provide a very dense and informative linkage marker set. However, it can be computationally intense to run such a linkage, with the bulk of the computation used in computing IBD. This IBD computation is the same for each analysis of different traits, so much computation time can be saved by computing the IBDs for the whole data set once.

We used Loki to estimate multipoint IBD with a sex-specific map. Loki uses MCMC estimation and can accurately estimate IBD in large pedigrees. An alternative would have been to use an implementation of the Lander-Green algorithm such as MERLIN, but that would have required splitting many of our pedigrees due to Lander-Green being exponential in family size. Also, MERLIN does not allow for a sex-specific map: the actual recombination rates differ between males and females, but often a sex-averaged is used as an approximation. Since the differences in the male and female maps can be large, we decided to use a sex-specific map in our IBD calculations.

While we think that our haplotype construction is very accurate, we recognize that no process is 100% perfect. To help control for both potential errors in the underlying genotyping and the haplotype construction process, we computed three sets of IBDs. For the first set of IBD’s, we used all the Haplotype markers, which we called the ALL set. For the second and third, we took two sets of every other marker (skipping over the missing bins) to produce an EVEN and ODD set of IBDs. In analysis, we use all three sets and, if there are no problems with the haplotype markers, the LOD plots should look the same. These IBDs have been used in the analysis of many traits in LLFS.

## Discussion

In all our data filtering, we focused on keeping as many subjects and markers as possible, while filtering out questionable data down to the individual call level. This results in a data set where we feel confident in the data provided, but in which there are some missing calls. We could have fewer missing calls if we excluded the subjects with depth of coverage between 20 and 30, where a large proportion of the missing calls are, but our examination of these subjects suggested that we could make some high-quality calls at the variant sites that had adequate coverage. Consequently, we made the choice to release data on these subjects even though their call rate may not be as high as subjects with higher depth of coverage. If the goal had been more complete data with fewer missing values, we would have filtered out more subjects and markers to get the same quality data.

One of the advantages we had in using family data is that we could use Mendel error rate as a metric to examine the effectiveness of nearly all our filters. Our specific Mendel error and lineage filters can only be applied in family data. However, some of the other filters such as the excess heterozygosity filter and the B-allele frequency filter, can be applied in any genomic DNA data set. We implemented the B-allele frequency tests out of a specific desire to address Mendel errors, but while we used reduction in Mendel errors as a specific validation for this filter, the actual application of the filter does not require family data.

We continue to refine our methods. Each time we get more data, we review what we have done previously and welcome suggestions for improved filters. We hope that our trials will be helpful to others as we have benefitted from the experiences of others.

## Acknowledgements

This work was supported by National Institute on Aging (grants U01AG023746, U01AG023712, U01AG023749, U01AG023755, U01AG023744, and U19 AG063893).

